# The genome of *Lactuca saligna*, a wild relative of lettuce, provides insight into non-host resistance to the downy mildew *Bremia lactucae*

**DOI:** 10.1101/2022.10.18.512484

**Authors:** Wei Xiong, Lidija Berke, Richard Michelmore, Dirk-Jan M. van Workum, Frank F.M. Becker, Elio Schijlen, Linda V. Bakker, Sander Peters, Rob van Treuren, Marieke Jeuken, Klaas Bouwmeester, M. Eric Schranz

## Abstract

*Lactuca saligna* L. is a wild relative of cultivated lettuce (*Lactuca sativa* L.), with which it is partially interfertile. Hybrid progeny suffer from hybrid incompatibilities (HI), resulting in reduced fertility and distorted transmission ratios. *Lactuca saligna* displays broad spectrum resistance against lettuce downy mildew caused by *Bremia lactucae* Regel and is considered a non-host species. This phenomenon of resistance in *L. saligna* is called non-host resistance (NHR). One possible mechanism behind this NHR is through the plant–pathogen interaction triggered by pathogen-recognition receptors, including nucleotide-binding leucin-rich repeats (NLRs) and receptor-like kinases (RLKs). We report a chromosome-level genome assembly of *L. saligna* (accession CGN05327), leading to the identification of two large paracentric inversions (>50 Mb) between *L. saligna* and *L. sativa*. Genome-wide searches delineated the major resistance clusters as regions enriched in *NLR*s and *RLK*s. Three of the enriched regions co-locate with previously identified NHR intervals. RNA-seq analysis of *Bremia* infected lettuce identified several differentially expressed *RLK*s in NHR regions. Three tandem wall-associated kinase-encoding genes (*WAK*s) in the NHR8 interval display particularly high expression changes at an early stage of infection. We propose *RLK*s as strong candidate(s) for determinants for the NHR phenotype of *L. saligna*.

## INTRODUCTION

Lettuce (*Lactuca sativa* L.) is a leafy vegetable grown in more than 100 countries, with a total yield of over 29 million tons in 2019 (FAOSTAT, 2019). One of the most important goals for lettuce breeding is the introgression of durable resistance against lettuce downy mildew, a destructive disease caused by the oomycete pathogen *Bremia lactucae* Regel (Lebeda et al., 2009). Outbreak of downy mildew disease leads to substantial yield and economic losses.

Wild relatives of lettuce are often used to introgress novel resistances (Lebeda et al., 2014; Parra et al., 2016). *L. saligna*, which belongs to the secondary gene pool of lettuce, is an important donor to enhance resistance to *B. lactucae* in cultivated lettuce (Netzer et al., 1976; Norwood et al., 1981; Bonnier et al., 1991). *L. saligna* is a diploid (2n=2x=18, same as lettuce) and self-pollinating species, which is partially interfertile with *L. sativa* (Lebeda et al., 2007, 2019). It is broadly distributed across Eurasia, from the Mediterranean region towards temperate Europe, and from the Iberian Peninsula to Central Asia (Zohary, 1991; Doležalová et al., 2002; Lebeda et al., 2019). *Lactuca saligna* is of particular interest to lettuce breeders as a potential resistance donor due to its complete resistance to all races of *B. lactucae.* As such, it is considered a non-host species to *B. lactucae* based on the definition: “All genotypes of a species are resistant against all genotypes of a specific pathogen” (Bonnier et al., 1991; Petrželová et al., 2011; Lebeda et al., 2009; Heath, 1981). For convenience, we term this resistance phenotype of *L. saligna* as non-host resistance (NHR), which is defined as strictly phenomenological and does not imply a molecular mechanism (Panstruga and Moscou, 2020). To successfully introgress this NHR in lettuce cultivars the gene(s) underlying the non-host resistance and the reproductive barriers observed in hybrid offspring should be determined.

Although *L. saligna* is crossable with *L. sativa*, the F_1_ plants are nearly sterile, and the resulting inbred offspring (F_2_ generation) show severely reduced fertility and transmission ratio distortions due to hybrid incompatibility (HI) (Jeuken et al., 2001; Giesbers et al., 2019). Some case of HI can be explained by the deleterious combination of interspecific alleles according the Dobzhansky-Muller (DM) model (Dobzhansky, 1934; Muller, 1942; Bateson, 1909). Many identified and resolved HI loci are explained by a digenic deleterious epistatic interaction and often results in transmission ratio distortion (TRD) (Fishman and Sweigart, 2018; Fishman and McIntosh, 2019). In F_2_ offspring and backcross inbred lines (BILs; i.e., single segment introgression lines) of *L. saligna* x *L. sativa*, 11 HI loci were associated with TRD, six of which were nullified by a paired allele from *L. saligna* (Giesbers et al., 2019). HI loci may reduce the efficiency of introgression of NHR genes from *L. saligna* into *L. sativa* when HI- and NHR loci are closely linked (i.e., linkage drag). In addition to HI, an interspecific chromosomal rearrangement, like an inversion, will also hamper the introgression of desired NHR genes via linkage drag caused by reduced recombination (Hoffmann and Rieseberg, 2008; Fishman and Sweigart, 2018).

NHR in plants is suggested to rely on a continuum of layered defenses, including both constitutive and induced resistance mechanisms (Niks and Marcel, 2009; Jones and Dangl, 2006; Bettgenhaeuser et al., 2014). Previous studies propose that induced NHR and host immunity rely on a similar non-self-recognition system comprising two innate immunity layers: i) pattern-triggered immunity (PTI) mediated by extracellular recognition of conserved non-self-molecules − called pathogen-associated molecular patterns (PAMPs) − by cell surface receptors, such as diverse receptor-like kinases (RLKs), and ii) host defense conferred by effector-triggered immunity (ETI) mediated by *R* genes encoding intracellular nucleotide-binding leucine-rich repeat proteins (NLRs) that recognize cognate pathogen-secreted effector molecules (Niks and Marcel, 2009; Schulze-Lefert and Panstruga, 2011; Jones and Dangl, 2006; Chisholm et al., 2006). After host penetration, hyphal growth of *B. lactucae* is quickly halted in *L. saligna* and consequently haustorium formation is impeded (Niks, 1987; Lebeda and Reinink, 1994; Zhang et al., 2009a, 2009b). To identify common loci associated with NHR to *B. lactucae* in lettuce, Giesbers *et al*. (2018) performed mapping studies based on nine *L. saligna* accessions from a broad range of geographic regions via multiple bidirectional backcrosses: i.e., i) BC1 populations in both parental directions (F_1_ x host *L. sativa*) and (F_1_ x non-host *L. saligna*), and ii) BC1S3 lines with three generations of inbreeding, respectively. These mapping populations facilitated the identification of four epistatic segments accounting for NHR in *L. saligna*: one positioned on Chromosome 4 (NHR4), two on Chromosome 7 (NHR7.1 & 7.2), and another on Chromosome 8 (NHR8) (Giesbers et al., 2018). It is worth noting that the NHR8 interval is closely linked to HI/TRD loci, which potentially limits fine-mapping and introgression of non-host traits into *L. sativa*. The genes and mechanisms underlying these four NHR loci are unresolved. A high-resolution analysis of these regions is needed to unveil the genetic determinants governing NHR in *L. saligna*.

*NLR*s in lettuce were previously identified by Christopoulou *et al*. (2015a) using the *L. sativa* v6 genome. Identified NLRs were classified into two major groups: TNLs with TOLL/interleukin-1 receptor (TIR) domains and CNLs with coiled-coil (CC) domains, and subsequently into multiple resistance gene candidates (RGC) families (Takken and Goverse, 2012; McHale et al., 2006; Meyers et al., 2003). Almost all identified *NLRs* were found to reside in five major resistance clusters (MRCs) that co-segregate with resistance to diverse pathogens (McHale et al., 2009; Christopoulou et al., 2015b). For example, MRC2 on Chromosome 2 comprises multiple *RGC2* family members, including the downy mildew resistance genes *Dm3*, *Dm14*, *Dm16*, and *Dm18* (Shen et al., 2002; Wroblewski et al., 2007; Christopoulou et al., 2015a). Similar MRCs are suggested to be present in *L. saligna* based on the expected whole-genome synteny, since some qualitative resistance phenotypes from *L. saligna* have been mapped at single loci syntenic to MRCs in *L. sativa* (Giesbers et al., 2017). An *L. saligna* genome reference can facilitate synteny analysis to recognize these anticipated MRCs.

Multiple *RLK* families contain members involved in a wide range of immune responses in plants. Notable examples can be found in the LRR-RLK sub-family, such as FLS2 involved in the perception of bacterial flagellin and IOS1 that contributes towards resistance to the downy mildew *Hyaloperonospora arabidopsidis* (Zipfel et al., 2004; Hok et al., 2011). Previously, Christopoulou *et al*. (2015a) also described *LRR-RLK* encoding genes in lettuce. Nevertheless, a specific inventory of genes encoding other resistance-related *RLK*s in lettuce, such as those encoding lectin receptor kinases (LecRKs) and wall-associated kinases (WAKs), is still lacking, not to mention the *RLK*s of *L. saligna* (Bouwmeester et al., 2011; Hu et al., 2017; Zuo et al., 2015; Hurni et al., 2015; He et al., 1999).

Here, we report a *de novo* genome assembly of *L. saligna* (accession CGN05327) using a variety of sequencing and scaffolding techniques. The assembly was compiled into nine chromosomal pseudo-molecules by genetic mapping. The resulting assembly enabled us to conduct diverse genomic analyses to dissect the genetic determinants underlying non-host resistance in *L. saligna*. The analyses provide insights of evolution into disease resistance and on host-pathogen arms race in lettuce. For breeding, the gained knowledge helps to facilitate the introgression of *Bremia* resistance into cultivated lettuce.

## RESULTS

### Genome sequencing and assembly

*L. saligna* accession CGN05327 was used to produce a reference genome for *L. saligna* (Supplemental Note). A combination of PacBio long-read (95.4 Gb; 41X) and Illumina short-read (407.4 Gb; 175X) sequencing was generated to assemble the genome (Supplemental Data 1). Illumina paired-end (125 bp, PE) and mate-pair (300 bp, MP) reads were generated from three libraries of different insert size (200 bp, 500 bp and 550 bp) (Supplemental Data 1A). The *L. saligna* genome size was estimated by K-mer analysis to be 2.27 Gb, which agrees with genome size estimates established by flow cytometry (2.3 Gb; Doležalová et al., 2002; Zohary, 1991). K-mer analysis also revealed that the genome is highly homozygous (estimated heterozygosity = 0.12%) as expected for this inbreeding species (Supplemental Figure 1 and Supplemental Table 1). To construct a high-quality genome of *L. saligna*, we applied a variety of advanced assembly and mapping techniques (Supplemental Figure 2). An initial Canu assembly (v0.5) consisted of 31,431 contigs, and the N50 number and size were 6,957 and 88.0 kb, respectively (Table 1; Supplemental Data 1A). Bionano fingerprinting, 10x Genomics barcoding, and Dovetail Hi-C library data were sequentially applied to construct the version 2 assembly, which refined the assembly to 24 super-scaffolds (largest scaffold = 279.9 Mb; N50 = 146.7 Mb; Supplemental Data 2B-C).

**Table 1.**
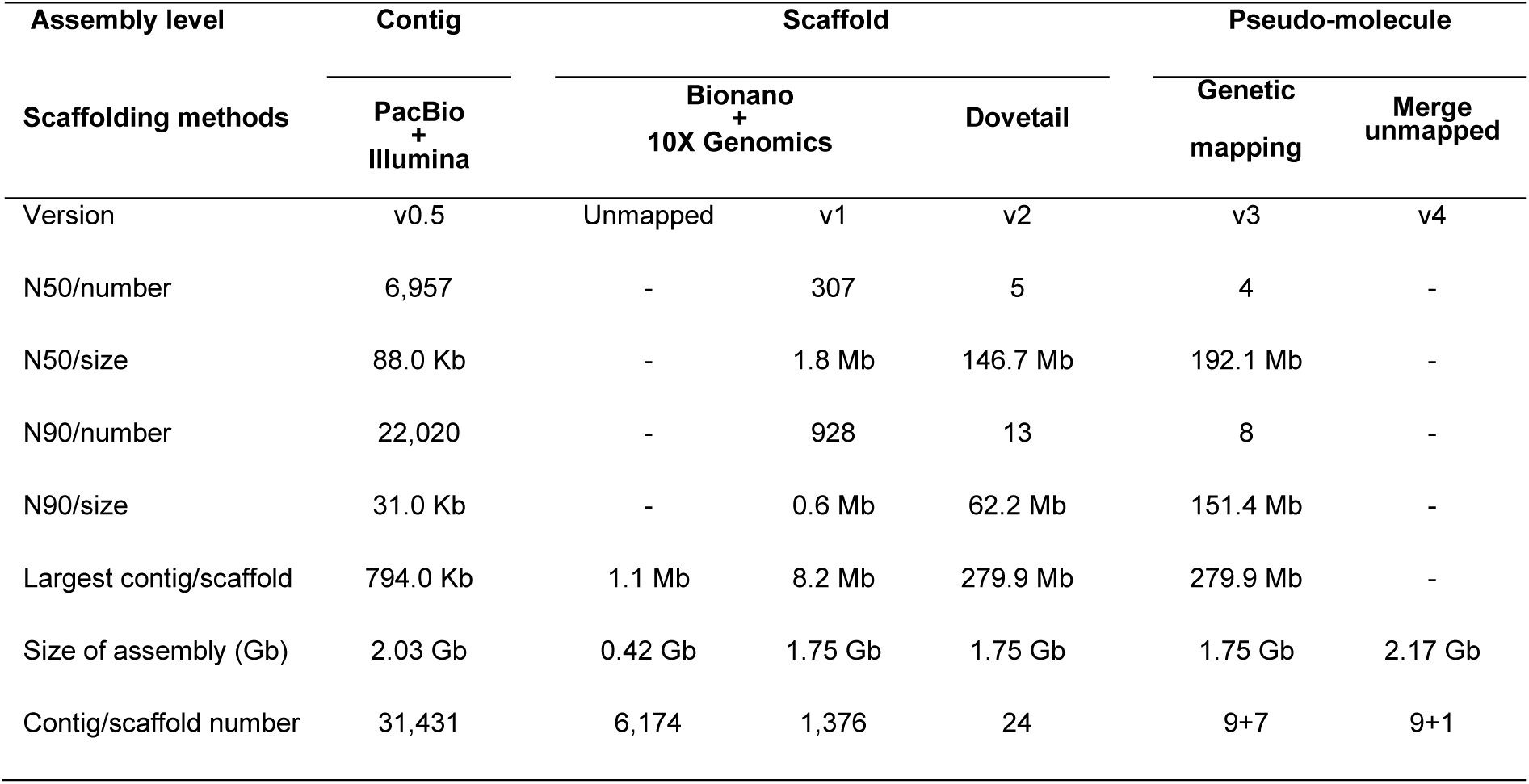
Genome assembly summary

### Linkage group anchoring and assembly assessment

To generate chromosomal pseudo-molecules, we combined 417 genetic markers from an F_2_ population linkage map (*L. saligna* x *L. sativa*) and 19,027 syntenic markers between *L. saligna* and *L. sativa* (Supplemental Table 2; Supplemental Data 3A-B). This resulted in a chromosome-level assembly (v3) in which 17 out of 24 scaffolds (99.8% bases, 1.75 Gb) were anchored and oriented into nine chromosomes, covering ∼77% of the estimated genomic sequence (1.75 of 2.27 Gb) (Supplemental Table 3; Supplemental Data 2D; Supplemental Figure 3). To obtain a more complete reference assembly, un-scaffolded contigs (>1000 bp) were merged to create a virtual “chromosome zero.” This eventually led to a final assembly (v4) with nine chromosomes plus chromosome zero, with a complete genome size of 2.17 Gb (Table 1; Supplemental Table 4; Supplemental Data 2E-F). This final assembly contains 91.9% (1,951 out of 2,121) of the expected BUSCO (1,859 single and 92 duplicated copies) eudicot gene models (Supplemental Table 5), and 92% of the 30,696 *L. saligna* expressed sequence tags (ESTs) in NCBI could be aligned to the v4 assembly at 80% identity and 80% coverage (Supplemental Table 6).

### Repeat and non-coding RNA annotation

Our analyses estimated that 77.5% of the *L. saligna* genome consists of transposable elements (TE; Table 2; Supplemental Table 7). Long terminal repeat retrotransposons (LTR-RT) were the most predominant repetitive elements, comprising both Gypsy and Copia retrotransposons (43.8% and 23.1% of genome, respectively) (Supplemental Table 8-9). TEs were distributed across the genome, and found to be enriched in regions roughly representing the pericentromeric locations (Supplemental Figure 4: track E). Non-coding RNAs involved in mRNA transcription (snRNA), translation (tRNAs and rRNAs), and regulation of gene expression (miRNAs) were also annotated (Table 2; Supplemental Table 10).

**Table 2.** Genome annotation summary

| Genome annotation | Metrics | Stats |
| --- | --- | --- |
| Gene prediction | *n* of genes | 42,908 |
|  | Mean length of CDS | 1,117 bp |
|  | Mean exon number | 5.1 |
|  | *n* protein-coding genes | 40,730 (94.9%) |
| ncRNA | *n* of rRNAs | 4,114 |
|  | *n* of tRNAs | 1,857 |
|  | *n* of miRNAs | 128 |
|  | *n* of snRNAs | 329 |
| Transposable elements | %Retrotransposons | 67.8% (1.5 Gb) |
|  | %DNA transposons | 3.1% (66.9 Mb) |
|  | %Unclassified repeats | 6.6% (143.6 Mb) |
|  | %Total | 77.5% (1.7 Gb) |

### Gene prediction and functional annotation

A combination of *de novo* search and homology support was applied for gene model prediction. Most of the predicted gene models (93%) were well supported (AED > 0.5) by RNA-seq data and gene homology (Supplemental Figure 5; Supplemental Data 4A). In total, 42,908 gene models were retained after filtering based on coding-potential (Table 2). The average coding size and exon number per gene was 1.3 kb and 5.1 respectively (Table 2). We further validated the potential for protein-encoding sequences using domain, ortholog, and homolog databases. By combining all results, 40,730 genes (94.9%) had matches in at least one database (Table 2; Supplemental Table 11; Supplemental Data 4B).

### *Lactuca saligna* population structure and diversity

To explore the genetic diversity and population structure of *L. saligna*, we re-sequenced 15 accessions representing the distribution across its native range (Supplemental Table 12-13). SNPs were first called on the *L. saligna* genome assembly and then filtered on missing rate (<10 %) and minor allele frequency (>0.05), yielding 5,170,479 SNPs for downstream analysis (Supplemental Table 14-15). After pruning the SNP dataset, we applied three complementary methods to explore the structure of *L. saligna*: neighbor-joining tree building, principal component analysis (PCA), and ancestry history inference. The neighbor-joining tree revealed that the *L. saligna* population can be subdivided into three major clades that are largely congruent with the geographical origins of the selected accessions (Figure 1A). This finding was recapitulated by PCA (Figure 1B) and ADMIXTURE analysis (Figure 1C). These analyses also uncovered the geographical origins of two accessions that were previously unknown. Accession CGN05271 is implicated to be of European origin, whereas CGN05282 groups with multiple accessions from the Middle East (Figure 1D). It is noteworthy to mention that accession CGN05271, now found to be of European origin, has been extensively utilized in many in-depth genetic studies on resistance to downy mildew or reproductive barriers (Jeuken and Lindhout, 2002; den Boer, 2014; Giesbers et al., 2017, 2018; Jeuken et al., 2001; Giesbers et al., 2019). Our sequenced reference CGN05327 is genetically clustered with CGN05271. Finally, the leaf morphology of each accession was also found in line with the *L. saligna* population genetic structure (Supplemental Figure 6).

**Figure 1.**
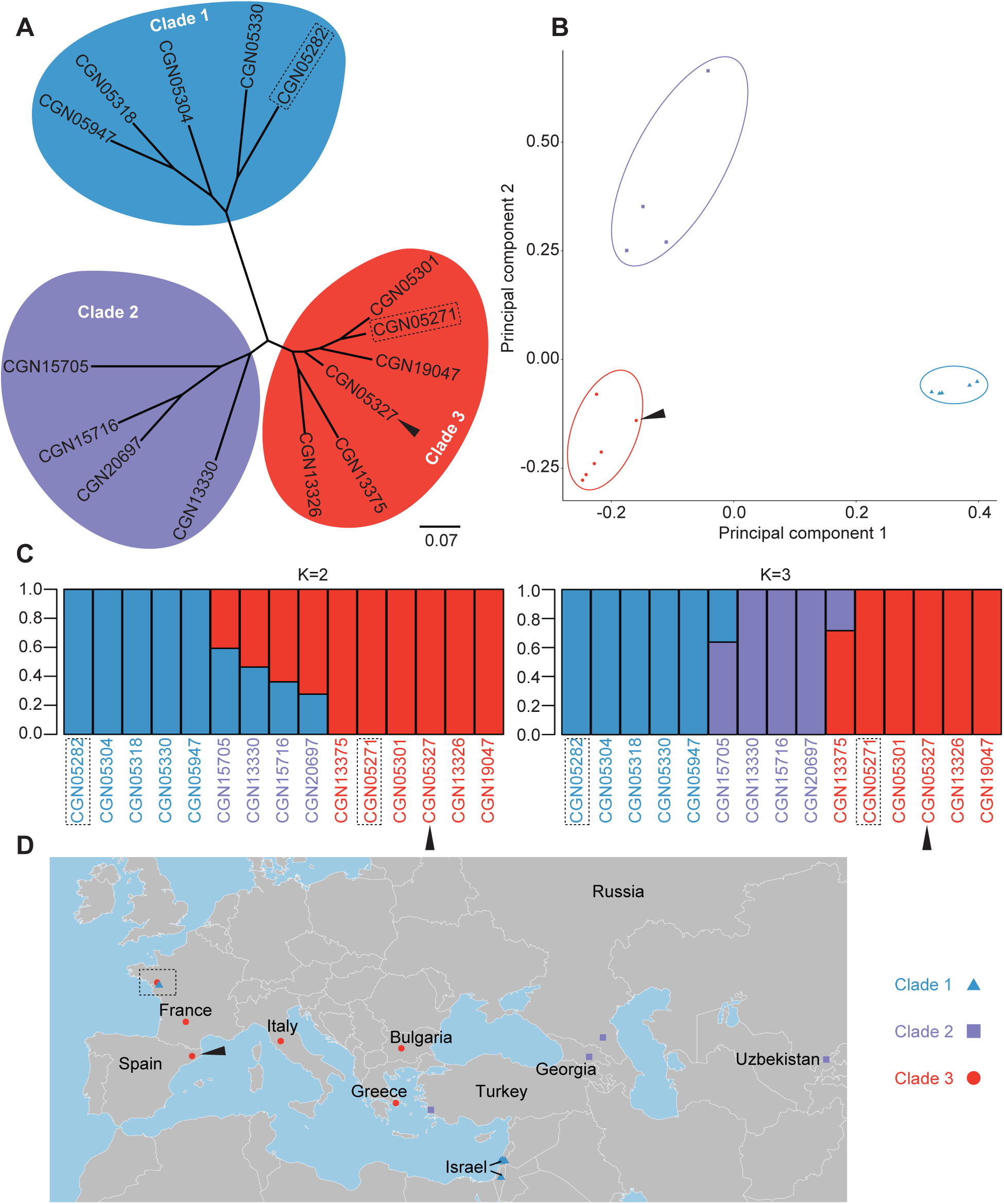
Resequencing of 15 accessions illustrates the *L. saligna* population structure. A, Neighbor- joining tree of 15 re-sequenced *L. saligna* accessions based on called SNPs. Accessions were clustered into three clades (colored in red, blue, and purple). Two accessions with unknown origins obtained from a French botanical garden are labelled by dashed lines. The black arrow indicates reference accession CGN05327 used for *de novo* sequencing. B, Principal component analysis plot of the top two- components illustrating the *L. saligna* population structure. Colors and shapes correspond to clades 1, 2, and 3. C, Genetic ancestry estimation with presumed populations (K=2 and K=3) indicating the population number and evolution. Red and blue represents the two ancestral populations and the purple indicates an intermediate population between the two ancestors. D, Geographic locations of *L. saligna* accessions, colored and shaped based on population structure.

### Synteny between L. saligna and L. sativa

Duplication events and structural variation were identified between the *L. saligna* and *L. sativa* genomes by syntenic alignments. Intra-species collinearity revealed a 3:1 syntenic pattern in all nine chromosomes for both species, confirming the known shared whole-genome triplication event within the Asteraceae (Reyes-Chin-Wo et al., 2017; Iorizzo et al., 2016) (Supplemental Figure 7). Inter-species syntenic analysis revealed a high level of genome-wide collinearity between both *Lactuca* species, except for two large inversions (> 50 Mb) on Chromosomes 5 and 8 (Figure 2B-C; Supplemental Table 16). The observed gene density (∼ 20 genes per Mb) within these two inverted regions in both species suggests that they are not close to the centromere, i.e., paracentric inversions (Supplemental Table 16). The ranges and positions of inversions were estimated using syntenic genes at the inversion borders (Supplemental Table 17). To confirm these inversions, we mapped markers derived from an interspecific F_2_ population to the *L. saligna* genome and compared their genetic and genomic positions. This showed that the genetic position plateaued while the genomic position kept increasing over the inverted region, which reflects the suppressed recombination due to inversion (Supplemental Figure 8). These inversions encompass of a diversity of genes, some of which encode proteins known to play key roles in various biological processes, such as a methyltransferase involved in Vitamin E biosynthesis and a phosphatase regulating cell wall integrity (Supplemental Table 18-19) (Cheng et al., 2003; Franck et al., 2018).

**Figure 2.**
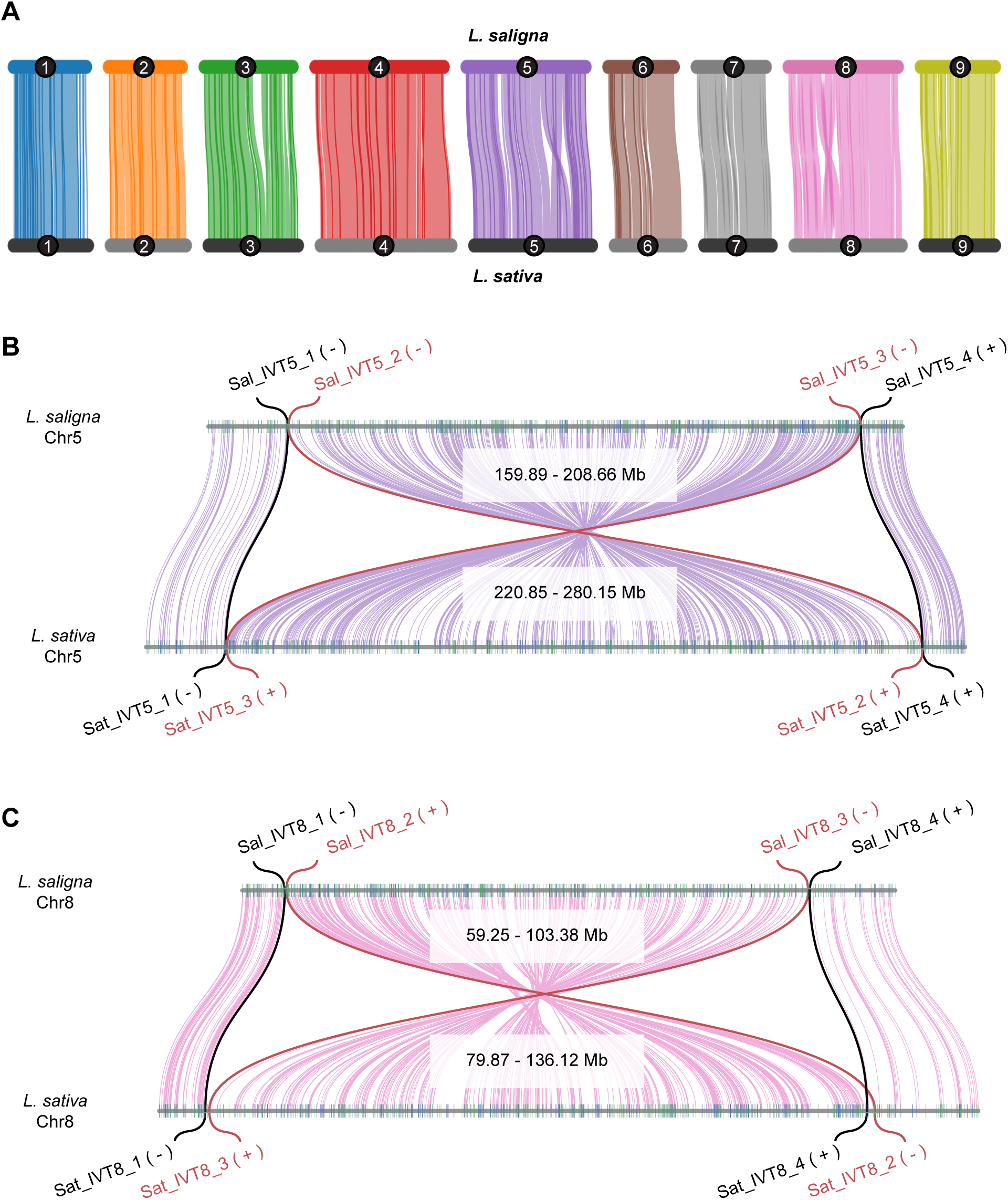
Synteny reveals two large inversions on chromosomes 5 and 8 between *L. saligna* and *L. sativa*. A, Synteny of best orthologs for each chromosome between the two *Lactuca* species. Each chromosome is represented by a different color. B-C, Inverted synteny regions on chromosome 5 (purple) and 8 (pink) with 50 flanking genes shown at borders, respectively. Relative to *L. saligna*, the red lines link the first and last homologous gene pairs within inverted synteny, while the black lines indicate the first homologous pairs outside of the inversion.

### Comparison of *NLR* content and distribution between *L. saligna* and *L. sativa*

To explore variation in the *NLR* gene family, HMMER and BLAST searches were conducted against the proteomes of *L. saligna* and *L. sativa* (Supplemental Data 5A- B). Retrieved amino acid sequences were first classified based on their N-terminal TIR or CC domain (TNLs and CNLs, respectively) and thereafter subdivided to Resistance Gene Candidate (RGC) families by phylogenetic analyses (Supplemental Figure 9; Supplemental Data 5C-D). This resulted in the identification of 323 *NLRs* in *L. saligna* and 364 *NLRs* in *L. sativa*. *Lactuca saligna* and *L. sativa* were found to contain a similar content of both TNL- and CNL-type, i.e., 184 versus 202 (57.0%, 55.5%), and 139 versus 162 (43.0%, 44.5%), respectively (Table 3; Supplemental Table 20). Genomic positions of MRCs previously identified in *L. sativa* were identified in the *L. saligna* genome assembly using *L. sativa* orthologs (Supplemental Table 21; Supplemental Data 5D). We additionally defined two *NLR*-enriched clusters (NCs) in *L. saligna* on Chromosomes 4 (38.55 – 40.68 Mb) and 7 (44.01 – 44.48 Mb), hereafter named NC4 and NC7 (Supplemental Table 22). These two NCs were also identified in *L. sativa,* but were not previously labeled as MRCs due to the absence of resistance phenotypes (Christopoulou et al., 2015b). In total, 41 RGC families were identified. Seven RGC families (six singletons and one multigene family) present in *L. sativa* were found missing in *L. saligna*, which might be caused by the reconstructed phylogeny or they may be unique to *L. sativa* (Supplemental Table 23). While *L. saligna* has a similar amount of NLRs compared to *L. sativa* in most RGCs, we defined significant size change by count and percentage difference. In this way, we observed that six and three RGC families were contracted (i.e. RGC1, 4, 8, 9, 14, and 21) and expanded (i.e. RGC 16, 20, and 29), respectively, in this accession of *L. saligna* compared to the reference genome of *L. sativa* (Supplemental Figure 9; Supplemental Table 23).

**Table 3.**
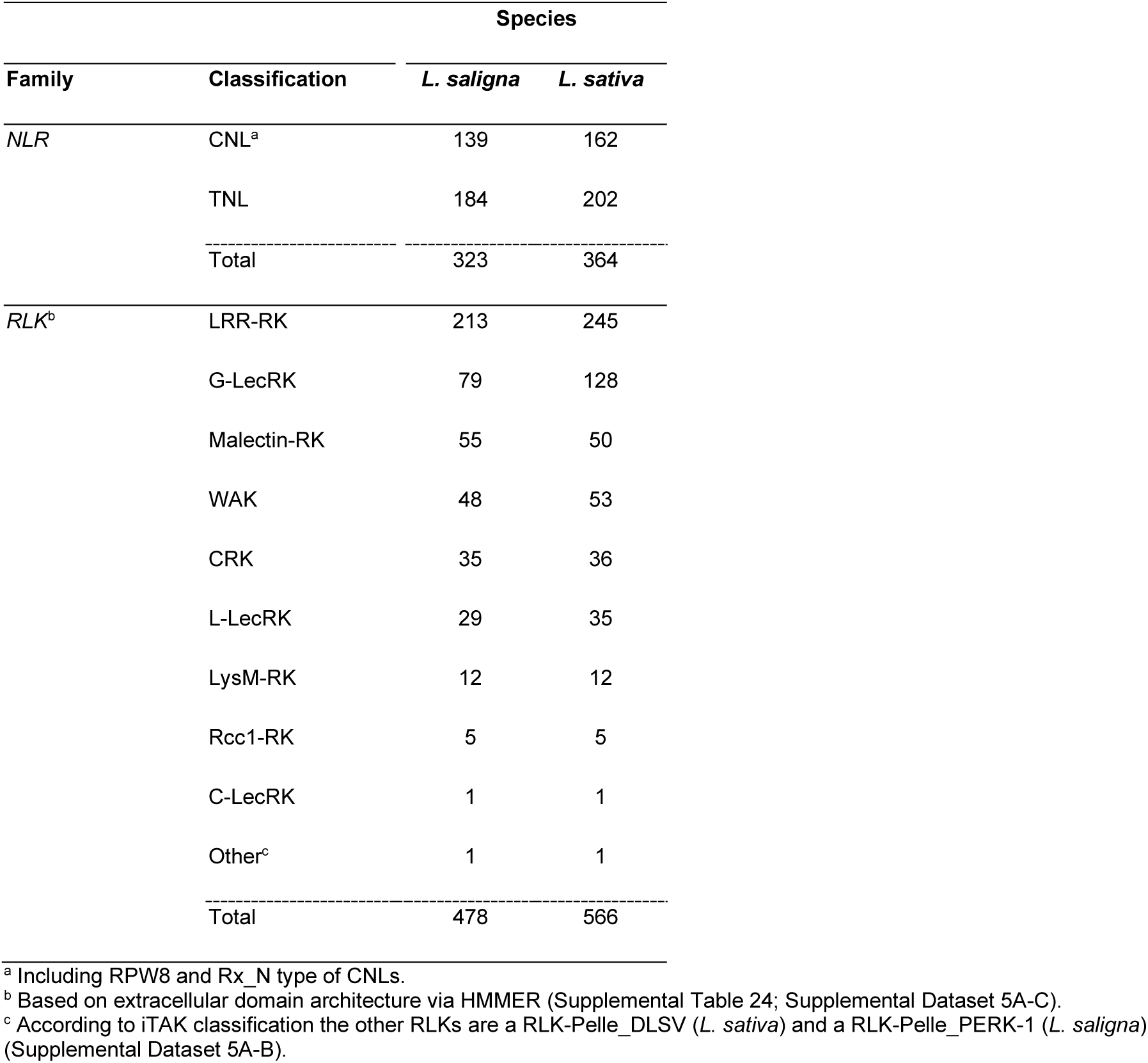
Identification and classification of *NLR*s and *RLK*s for *L. saligna and L. sativa*

### Comparison of *RLK* genes between *L. saligna* and *L. sativa*

To identify genes encoding RLK proteins, we performed in-depth HMMER searches against the predicted *L. saligna* and *L. sativa* proteomes. This resulted in the identification of 478 and 566 *RLK* encoding genes in *L. saligna* and *L. sativa,* respectively (Supplemental Table 24; Supplemental Data 6A). Sliding window analysis revealed that *RLKs* are distributed on all chromosomes, with their density elevated at the chromosomal ends (Figure 3: track B). RLKs were further classified into nine subfamilies based on their extracellular domains using HMMER (Table 3; Supplemental Data 6B). In both species, LRR-RLKs and G-type LecRKs (G-LecRKs) formed the largest subfamilies. The major difference in total *RLK*s was also largely accounted by these two subfamilies − with an additional 32 G-LecRKs and 48 LRR-RLKs in *L. sativa*. The other *RLK* subfamilies were found to be of similar size in these accessions of both *L. saligna* and *L. sativa*.

**Figure 3.**
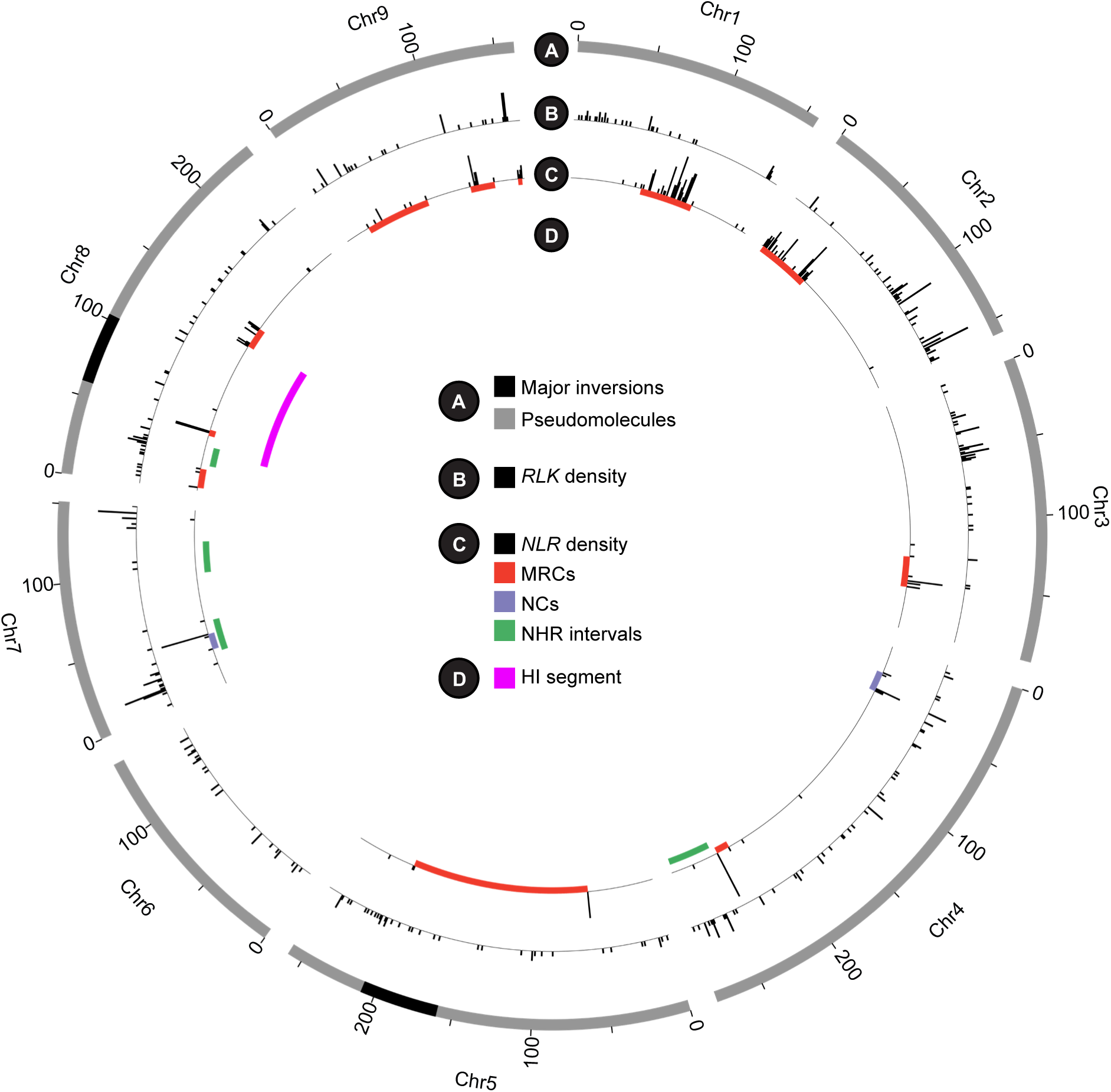
Phenotype mapping associates immune gene hotspots with NHR and HI regions. Track A, Circular ideogram of the nine pseudo-chromosomes (Mb) of the *L. saligna* assembly indicating two major inverted regions between *L. saligna* and *L. sativa.* Track B, Histogram of *RLK* density (1Mb window). Track C, *NLR* density (1Mb window) and tiles related to disease-resistance gene cluster intervals: i.e. major resistance clusters (MRCs), *NLR* clusters (NCs) with elevated density, and previously identified NHR interval. Track D, HI segment found on chromosome 8 using backcross inbred lines (*L. saligna* x *L. sativa*).

### Mapping HI and NHR loci on *L. saligna* genome

To precisely characterize the HI and NHR regions, markers of these loci were mapped to the *L. saligna* assembly (Supplemental Table 25-26). For HI, one locus was positioned on Chromosome 8 (33.15–138.07 Mb) and contains the inversion identified on Chromosome 8, 59–103 Mb (Figure 3: track A and D; Supplemental Figure 8) (Giesbers et al., 2019). This HI region and inversion region on Chromosome 8 was also adjacent to the resistance-related regions NHR8, MRC8B, and MRC8C (Figure 3: track C-D; Supplemental Figure 8). For NHR, three out of four intervals either overlapped with *NLR* or *RLK* hotspots. NHR7.1 was found to co-segregate with the NC7 region encoding 13 *NLR*s, whereas the other three NHR intervals consist of no or only one *NLR* gene (Supplemental Data 7). Moreover, mapping revealed that both NHR4 and NHR8 co-locate with regions enriched in *RLK* genes (NHR4: 20 *RLKs* in 34.21 Mb; NHR8: 14 *RLKs* in 13.26 Mb). Especially for region NHR8, the *RLK* density (1.06/Mb) was five-times higher than the genome-wide average (0.24/Mb, 422 in 1,745 Mb, excluding Chromosome 0). Mapping revealed a close relationship between NHR regions and resistance gene hotspots, making NLRs/RLKs potential determinants of NHR in *L. saligna*. In addition, NHR8 is also positioned near an HI segment, which may prevent the introgression of the candidate resistance genes to cultivated lettuce, impacting breeding for resistance.

### RNA-seq time-course analysis of *L. saligna* transcriptome in response to *Bremia*

To detect genes with differential expression after infection, we performed a *Bremia* infection assay on leaves of *L. saligna* to generate transcriptomic data and subsequently conducted a differential expression (DE) analysis. Treated and control samples were collected at 8- and 24-hours post-infection (hpi). Statistical analysis of quantified RNA-seq reads count identified a total of 1,268 and 1,688 differentially expressed genes (DEGs) (padj < 0.5 and log2FC > 1) at 8 hpi and 24 hpi, respectively (Supplemental Table 27; Supplemental Data 7). For both time points, the majority of DEGs were up-regulated in expression, i.e. 1,222 up-regulated versus 46 down- regulated genes at 8 hpi, and 1,362 up-regulated versus 326 down-regulated genes at 24 hpi (Supplemental Table 28). One of the most representative DEGs is Lsal_1_v1_gn_1_00001954, showing the largest induction in expression (log2FC=11.72), is a homolog of the penetration resistance gene *PEN1*, which encodes a syntaxin involved in vesicle assembly for non-host resistance against powdery mildew penetration in *Arabidopsis* (Collins et al., 2003).

### Enrichment analysis of identified DEGs in *L. saligna*

Subsequently, we applied gene ontology enrichment analysis of DEGs to explore functional-related biological processes and pathways. Figure 4 shows the 20 most significantly enriched terms related to DEGs at 8 hpi or 24 hpi. Sixteen out of 20 ontology terms were identified at both time points. Most clusters were mainly associated with resistance responses, like stress perception (GO:0009620), signal transduction (GO:0046777), and cell death (GO:0008219). In general, 8 hpi showed a greater enrichment than 24 hpi for most top terms (Figure 4B). In contrast, three unique biological clusters were found for the 24 hpi timepoint, all of which were related to ribosome biogenesis (GO:0042254, GO:0042273, and ath03010) (Figure 4A-B). In addition to the top 20 terms, many up-regulated genes were found to be involved in plant defense, in particular in response to oomycetes, illustrating the immune response of *L. saligna* upon *Bremia* infection (Supplemental Figure 10; Supplemental Table 28-29). For example, these include Lsal_1_v1_gn_9_00004094, a homolog of the lectin receptor gene *LecRK-IX.1* conferring resistance to *Phytophthora* spp. (another oomycete pathogen); Lsal_1_v1_gn_8_00004656 (*SARD1*) and Lsal_1_v1_gn_2_00003439 (*UGT76B1*), encoding two key regulators of salicylic acid (SA) synthesis and SA mediated signaling for stress response (Wang et al., 2015; Ding et al., 2016; Mohnike et al., 2021; Bauer et al., 2021). Our enrichment analysis detected that DEGs at both time points post-inoculation with *Bremia* were enriched in resistance-related biological processes: 8 hpi showed a stronger signal of early immune response and 24 hpi showed a shift of enriched terms to extra post-transcriptional response.

**Figure 4.**
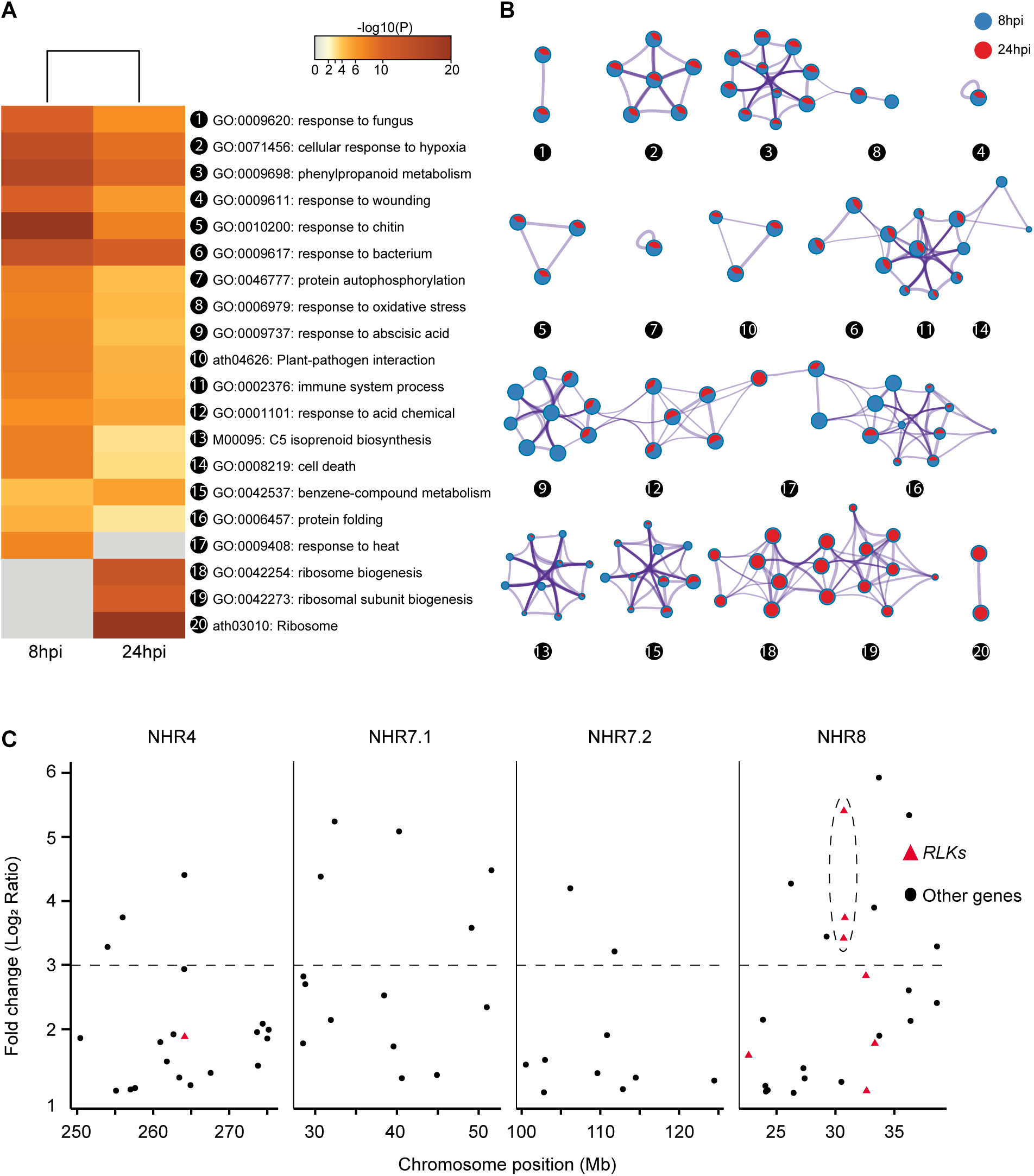
Enrichment analysis and expression levels of DEGs in *L. saligna* upon *Bremia* infection. A, Heatmap of the top 20 ontology groups at 8 and 24 hpi. Each group comprises multiple ontology terms and is represented by the term with the best p-value. Groups are hierarchically clustered and heatmap cells are colored according transformed p-values [-log10(p-value)]. Grey cells indicate a lack of enrichment for that term in the corresponding gene list. B, Networks of representative terms for the top 20 groups. Each term is displayed by a pie chart node to illustrate the proportional number of up- regulated genes at 8 hpi (blue) and 24 hpi (red). Some groups are interconnected and form a larger network. C, Distribution of up-regulated genes across the four identified NHR regions in *L. saligna* at 8 hpi. The horizontal dashed line (y=3) indicates the cutoff for up-regulated genes (log2FC > 3). Receptor-like kinases (*RLK*s) are indicated by red triangles, and other genes are black circles. The dashed ellipse line points out the three tandem arrayed *WAK*s on NHR8.

### Differentially expressed genes in NHR regions at 8 hpi

Based on above mapping and DE analysis results, we inspected the statistics of the up-regulated genes in NHR intervals at 8 hpi to further identify candidates for resistance to lettuce downy mildew. First, we calculated the DEG density per million base-pair of four NHR loci and the whole genome (Supplemental Table 30). As baseline, the DEG density for the whole genome was 0.70 per Mb. The NHR8 locus had the highest DEG density (1.54/Mb) among all NHR intervals and was greater than two-times the average density of the entire genome. Moreover, 11 DEGs located in the overlapping region of NHR8 and HI may also inhibit the ability to overcome the hybrid barrier. Secondly, we examined the percentage of up-regulated *RLKs* and *NLRs* (up- regulated number / total number) for each NHR interval (Supplemental Table 30). The percentage of differentially expressed *NLRs* was low (4.6%) across the whole genome. None of the *NLR*s within the two NHR loci were differential expressed. In contrast, more than 22.7% of the *RLKs* (96) were up-regulated genome-wide after *Bremia* inoculation. NHR8 also displayed a high percentage of up-regulated *RLKs* (50%, seven out of 14). Furthermore, we counted the number of DEGs with a large degree of change (log2FC > 3) in NHR regions of interest (Supplemental Table 30). Again, NHR8 was found to contain more highly expressed genes (nine) than the other three NHR regions. Thus, out of four NHR loci, the statistics of DEGs strongly suggests that genes on NHR8 seemed to play a critical role in the resistance to *Bremia*, especially the *RLK*s. Based on these observations, we pinpointed eight DEGs located in NHR8 as candidates for downy mildew resistance in *L. saligna* (Supplemental Table 31). One of the candidate genes encodes a plant U-box type E3 ubiquitin ligase (PUB), of which family members have been reported to play essential roles in plant defense and disease resistance (González-Lamothe et al., 2006). The other candidate genes all encode receptor-like kinases, i.e., one LysM-containing receptor-like kinase (LysM-RK), three G-type lectin receptor kinases (G-LecRKs), and three WAKs. It is noteworthy to mention that the three *WAK*s were tandem-arrayed, of which two were highly up-regulated (log2FC > 3; Figure 4C).

## DISCUSSION

### *L. saligna* reference genome and population structure

In this study, we report on the *de novo* genome assembly of *L. saligna* based on long- and short-read sequencing together with advanced scaffolding techniques. The genome size of our *L. saligna* assembly (2.17Gb) is in line with the previously reported C-value (2.3Gb) (Doležalová et al., 2002). The genomic content, such as gene space and repeat content of the genome (∼77%), is comparable to cultivated lettuce (Reyes-Chin-Wo et al., 2017). Using SNPs called on the reference genome, population genetic analysis identified three *L. saligna* sub-groups that are consistent with geography Figure 1). We also inferred the graphical origin of two genotypes derived from the Jardin Botanique de Nantes, a French botanical garden, including the accession CGN05271 (found to be of European origin) and accession CGN05282 (found to be of Middle Eastern origin). The obtained population genetics structure is in agreement with a previous clustering based on AFLP markers (Giesbers et al., 2018).

### Inversions and HI may hamper breeding with the NHR8 resistance locus

Comparative genomic analysis identified two large inversions (>50 Mb) on Chromosomes 5 and 8 between *L. saligna* and *L. sativa* (Figure 2). We also found that the inversion on Chromosome 8 co-segregated with an HI region (Giesbers et al., 2019). Genic incompatibilities associated with hybrid necrosis are often linked to immune genes (Bomblies and Weigel, 2007; Fishman and Sweigart, 2018). A well-described example of hybrid necrosis for lettuce is the digenic interaction between the *L. saligna* allele of *Rin4*, encoding a putative negative regulator of basal plant defense, and the resistance gene *Dm*39 from cultivated lettuce (Jeuken et al., 2009). The HI locus on Chromosome 8 was not found to be associated with the hybrid necrosis phenotype (Giesbers et al., 2019). Therefore, immune gene(s) are likely not causal to HI on Chromosome 8, even though we found several resistance loci (MRC and NHR) close to the HI regions located on the inverted regions (Figure 3; Supplemental Figure 8). If the HI/TRD locus indeed resides in the inversion, then further fine mapping and introgression of loci associated with HI, and potential immune genes underlying NHR for that matter, will not be feasible due to the lack of recombination caused by inversion. Future research could investigate whether all *L. saligna* accessions share same large inversions that cause linkage drag, by combining sequencing.

### *L. sativa* contains more immune genes than the non-host *L. saligna*

Previous studies in *L. saligna* by genetic mapping have detected multiple loci containing *NLR*s associated with its resistance phenotype, for example, the R locus (*Dm39*) interacts with *Rin4*, and the R locus responds to the effector *BLR31,* which are suggested not to govern the NHR phenotype (Jeuken et al., 2009; Giesbers et al., 2017). The lack of knowledge on genome-wide variation in resistance genes has hindered the identification of NHR determinant(s). In this paper, we comprehensively inventoried *NLR* and *RLK* genes in *L. saligna* and *L. sativa* (Table 3). Our results show that *L. sativa* has more *NLRs* (364 / 323 = 1.13) and *RLK*s (566 / 478 = 1.18) than *L. saligna*. This difference could possibly be due to the genome size differences between *L. sativa* and *L. saligna* (2.5Gb / 2.3Gb = 1.09; Doležalová et al., 2002) or the incomplete sequencing and annotation. Immune genes, like *NLRs* or *RLK*s, are known as the most variable genes in plants, including lettuce and its wild relatives (Karasov et al., 2014; Parra et al., 2016). Due to allelic and copy number variation, the genome assembly alone cannot fully capture the complete spectrum of *R* genes (Barragan and Weigel, 2021). Therefore, the genetic determinant of NHR might not be identified by the genome-wide searches using these reference assemblies. Target sequencing of *NLR*s and *RLK*s (e.g. RenSeq and RLKSeq) can be applied to collect a more complete spectrum of resistance genes (Witek et al., 2016; Lin et al., 2020).

### *RLK* and *NLR* genes associated with NHR against *Bremia*

To further understand the relationship between *RLK*/*NLRs* and NHR in *L. saligna*, we mapped the four NHR loci to the *L. saligna* reference genome. Of the four NHR intervals, we found that three have either elevated densities of *RLKs* (NHR4 & NHR8) or *NLR*s (NHR7.1). Moreover, *RLKs* and *NLRs* do not co-occur with each other in analyzed *Lactuca* species, as illustrated by NHR8 (14 *RLKs* vs zero *NLRs*) and NHR7.1 (zero *RLK*s vs 13 *NLRs*) (Supplemental Table 30), which suggests that *NLR*s and *RLK*s act as epistatic genes explaining NHR (Giesbers et al., 2018). Although *RLKs* and *NLRs* elicit PTI and ETI respectively (Jones and Dangl, 2006), there is increasing evidence that PTI and ETI are not separate phenomena and mutually strengthen each other’s immune response (Yuan et al., 2021; Ngou et al., 2021). This could explain why the identified NHR loci in *L. saligna* involves a combination of PTI and ETI.

### RNA-seq highlights a crucial role of *RLKs*

RNA-seq analysis of *L. saligna* leaves inoculated with *Bremia* enabled us to identify DEGs related to NHR-associated plant defense responses. Multiple DEGs with high levels of induced expression were found to be involved in salicylic acid (SA) synthesis (SARD1 and UGT76B1) or SA-dependent penetration resistance (PEN1 and PEN3) contributing to NHR in *Arabidopsis* (Supplemental Table 27-28) (Zhang et al., 2010; Collins et al., 2003; Assaad et al., 2004; Mohnike et al., 2021; Bauer et al., 2021). Various studies have shown that SA increases *RLK* expression in different plants (Ohtake et al., 2000; Coqueiro et al., 2015). Transcriptome analysis also revealed that a large portion of the DEGs at 8 hpi function in early recognition and defense signaling activity, whereas DEGs at 24 hpi were found to be responsible for post-transcription activity. For expression of immune genes in NHR regions, no *NLR* genes were differentially expressed. This is consistent with expectations, as NLR-encoding genes generally are lowly expressed after *Bremia* infection (Wroblewski et al., 2007). In addition, a large amount of *RLK*s was differentially transcribed, which is similar as described for the interaction between lettuce and the fungal pathogen *Botrytis cinerea* (De Cremer et al., 2013).

### NHR8 contains *WAK* genes highly upregulated upon *Bremia* infection

Among the four NHR regions, NHR8 has the highest number of differentially expressed *RLK*s (Supplemental Table 30). Within it, three closely clustered wall-associated kinases (WAKs) were of special interest because of their significant expression change (Figure 4C). These three WAK paralogs were homologs of *Arabidopsis WAK2,* which is highly expressed in leaves and can be up-regulated in expression upon pathogen infection and SA application (He et al., 1999). Various studies illustrated that WAKs provide quantitative resistance against various diseases in crops such as maize and rice (Zuo et al., 2015; Hurni et al., 2015; Hu et al., 2017). For *L. saligna* infected by *Bremia*, oligogalacturonides derived from damaged cell walls could be perceived by WAKs to trigger PTI (Raaymakers and Van den Ackerveken, 2016; Ferrari et al., 2013; Brutus et al., 2010). WAKs have also been implied in cell wall reinforcement. In rice, Xa4 strengthens the cell wall by promoting cellulose synthesis and suppressing cell wall loosening, thereby enhancing resistance to bacterial infection by *Xanthomonas oryzae* (Hu et al., 2017). Hence, WAKs located on NHR8 seem to hold potential in *L. saligna* resistance. Nevertheless, we cannot rule out the possibility that other genes/factors play roles in NHR in *L. saligna*, and the expressions level of *WAK*s along with other genes mentioned in this paper need to be further compared to their homologs in susceptible lettuce cultivars or resistant introgression lines. Future fine mapping and knock down/out experiments are needed to further pinpoint key factors underlying the NHR in *L. saligna* using the reference genome assembly presented in this paper.

### A model for NHR in *L. saligna* against lettuce downy mildew

Based on our findings and previous research, we propose an NHR model for *L. saligna* with the following three elements: i) The host status of *L. sativa* and *L. saligna* is partly determined by the variation in orthologous RLKs involved in immunity. A specific ortholog in *L. saligna* can effectively enhance resistance to colonization by *B. lactucae*. A comparable role of orthologous RLKs has been observed in the interaction between barley and leaf rust fungi, in which a LecRK of wild barley quantitatively enhances resistance (Wang et al., 2019). ii) After non-self-recognition by RLKs, cell wall-plasma membrane interactions are strengthened (Wolf, 2017), restricting intercellular hyphal growth. This is in line with the reduced hyphae formation found in infected *L. saligna* (Zhang et al., 2009b). In case of successful penetration, NHR to powdery mildew in barley is often backed up by NLR-mediated hypersensitive response (HR) (reviewed in Niks and Marcel, 2009). As for the observed NHR in *L. saligna*, this might also be the case.

## MATERIALS AND METHODS

### Plant materials and DNA isolation

*L. saligna* accessions selected for whole-genome sequencing and resequencing were obtained from the lettuce germplasm collection of the Centre for Genetic Resources, The Netherlands (CGN) (Supplemental Table 12). Accession CGN05327 was collected from Gerona, Spain, of which a Single Seed Descendant (SSD) was used for *de novo* reference genome sequencing and assembly. Re-sequencing data of 15 Single Seed Decent (SSD) lines derived from *L. saligna* accessions (Supplemental Table 12) were selected to represent the *L. saligna* germplasm. Seeds were stratified at 4°C for three days to improve germination. Seedlings were subsequently grown in a growth chamber at 17–19°C with LED light under a 16 h photoperiod and a relative humidity of 75–78%. After eight weeks, plants were transplanted to larger pots containing potting soil and grown under greenhouse conditions. Images of leaves (third mature leaf counted from the base) of 10 accessions belonging to different subgroups were taken from 15-week-old plants, which were grown in triplicate (Supplemental Table 32). Tissue sampling was performed when plants were close to bolting, and DNA was extracted using the protocol as in Ferguson *et al*. (2020).

### Genome sequencing

A *de novo* genome assembly of *L. saligna* CGN05327 (Supplemental Figure 1) was assembled using a ∼21-fold coverage of long-read data generated by PacBio Sequel technology (4,083,751 reads; N50 read length=16,581 bp; subread length=8,514 bp), and a ∼175-fold coverage of Paired-end (PE) reads obtained by Illumina mate pair sequencing. The mate pair library was prepared using different insert sizes and read lengths: HiSeq (200 bp insert size, 125 bp PE), HiSeq (500 bp insert size, 125 bp PE), and MiSeq (550 bp insert size, 300 bp PE).

### Genome assembling and scaffolding

PacBio reads were assembled using Canu and polished with Pilon (v1.20) using Illumina data (Koren et al., 2017; Walker et al., 2014). Subsequently, multiple techniques were applied to elevate the contiguity of the assembly. A follow-up assembly (version 1) was scaffolded using 10x Genomics Chromium barcoding data (ARC pipeline) and ∼130-fold coverage BioNano optical mapping data (Yeo et al., 2017). A Hi-C library produced by Dovetail Genomics providing ∼2,553-fold coverage of sequence data (429 million 2x150 bp read pairs) was used for *in vitro* proximity ligation. Mis-joins in assembled contigs were corrected using the HiRise pipeline, resulting into genome assembly v2 (Putnam et al., 2016).

### Assembly reconstruction by syntenic and genetic makers

ALLMAPS was applied to reconstruct scaffolds of the *L. saligna* v2 assembly to chromosomal linkage groups using two types of markers: 417 genetic markers (weight = 2) derived from F_2_ (*L. saligna* CGN05271 x *L. sativa* cv. Olof), and syntenic markers (weight = 1) derived from the reciprocal best hits between *L. saligna* v2 and *L. sativa* v8 (Jeuken et al., 2001; Giesbers et al., 2019; Tang et al., 2015b). Contigs (>1 kb) not clustered in chromosomes were concatenated by JCVI with 100 N-content gaps to generate a virtual “chromosome zero” storing left genetic content (Tang et al., 2015a).

### Genome size estimation

Two paired-end Illumina libraries of *L. saligna* were used for genome size estimation (∼117 Gb pairs; ∼932 million reads) using a k-mer count size of 23 (Supplemental Table 1). Jellyfish v2.3.0 was used to count the k-mer frequency (Marçais and Kingsford, 2011). Jellyfish output was used by GenomeScope (v2.0) to estimate haploid genome length, percentage of repetitive DNA, and heterozygosity of the *L. saligna* genome using the histogram file (Vurture et al., 2017).

### Genome completeness assessment

Completeness of the *L. saligna* genome assembly was evaluated using multiple approaches. BUSCO (v3.0.2) assessment was conducted using the eudicotyledons_odb10 database (Simão et al., 2015). In addition, 226,910 ESTs of diverse *Lactuca* species (retrieved on July 2019 by NCBI) were aligned to the genome using GMAP (version 2019-06-10) (Wu and Watanabe, 2005). For GMAP alignment, presence/absence of ESTs was determined after filtering the alignments by identity and coverage at different levels of stringency using custom scripts.

### Repeat annotation

Tandem Repeats Finder v4.04 was used to detect tandem repeats using the following parameters: Match=2, Mismatch=7, Delta= 7, PM=80, PI=10, Minscore=50, and MaxPeriod=2000 (Benson, 1999). TEs were searched using RepeatMasker v4.0.7 against ortholog and *de novo* databases in a serial order (Smit et al., 2019): i.e., by using orthology data from Repbase and Dfam (version 20170127), and *de novo* TEs library generated by RepeatModeler v2.0 and MITE-Hunter (Han and Wessler, 2010; Jurka et al., 2005; Hubley et al., 2015; Price et al., 2005). Perl tool “One code to find them all” was used to parse and quantify the number and position of predicted repeat elements (Bailly-Bechet et al., 2014).

### Non-coding RNA annotation

Non-coding RNA (ncRNA) loci were annotated according to different types. tRNAscan-SE v2.0.4 was used to annotate tRNAs using eukaryote parameters (Lowe and Eddy, 1997). In addition, rRNA was annotated using RNAmmer v1.2 (Lagesen et al., 2007). INFERNAL v1.1.2 was used to search against the Rfam database (release 14.1) to detect additional miRNA, snRNA, tRNA, rRNA, and snoRNA sequences (Kalvari et al., 2018; Nawrocki and Eddy, 2013). Annotations predicted by different tools were merged and condensed using GenomicRanges in R v3.6 (Lawrence et al., 2013).

### Infection assays

Leaves of three-week old *L. saligna* plants (accession CGN05327) were spray-inoculated with a spore suspension of *B. lactucae* race Bl:21 (2.0*10^5^ conidiospores/mL) or with sterile water. Treated plants were first kept in the dark for 4 h to maximize spore germination, and then incubated in a growth chamber at 15°C and a 16/8 h (day/night) photoperiod. Leaf samples of inoculated and mock-treated leaves were collected at 8 and 24 hpi. Leaf samples from the three biological replicates were immediately frozen in liquid nitrogen and stored in -80°C until further use.

### RNA library preparation and sequencing

Total RNA was isolated from 12 infection assay samples and one pooled sample consisting of root and flower bud material (pooled from different floral stages) using a Direct Zol RNA Miniprep Plus kit (Zymo Research) followed by DNAse treatment. RNA was purified by ethanol precipitation. Concentration and purity of RNA samples was measured with a Nanodrop 2000c spectrophotometer and a Qubit 4.0 fluorometer using a RNA Broad Range assay (Thermo Fisher Scientific). Paired-End sequencing (2 x 125 bp) was performed on an Illumina HiSeq2500 platform using two flow cell lanes.

### Gene prediction

Gene models for protein-coding genes were annotated by combining *ab initio* prediction and homology-based annotation. First, BRAKER was used to train an Augustus model with RNA-seq data to predict genes *ab initio* (Hoff et al., 2016). Thereafter, MAKER was applied to integrate the *ab initio* prediction with extrinsic evidence: i.e., *de novo* transcripts assembled by Trinity and protein homology data (Holt and Yandell, 2011; Grabherr et al., 2011). Annotation-edit-distance (AED) calculated by MAKER was used to examine the quality of the genome annotation. Coding-potential was calculated by CPC2 (Kang et al., 2017) to further filter out non-coding transcripts (Supplemental Data 4).

### Functional annotation

Potential biological function of proteins was inferred using three criteria: i) best-hit matches in SwissProt, TrEMBL, and *A. thaliana* Araport11 databases using BLAST v2.2.31 and DIAMOND (Buchfink et al., 2015) (E-value cut-off = 1e-5); ii) protein domains/motifs identified by InterProscan against the Pfam protein database (El-Gebali et al., 2018; Zdobnov and Apweiler, 2001); and iii) gene ontology (GO) based on InterPro entries. Orthology searches for pathway analysis were conducted with Kofamscan (Aramaki et al., 2019) using a customized HMM database of KEGG Orthologs (Kanehisa, 2000).

### Resequencing and SNPs calling

Libraries of PE reads (2 x 150 bp, insert size distribution peaks at 190 bp) were constructed and sequenced. Re-sequencing reads were mapped to the *de novo* genome assembly using the BWA alignment tool (Li, 2013). After mapping, the alignment output in SAM format was translated to the BAM format using SAMtools (Li et al., 2009). Duplicated reads were marked, and read groups were assigned to the remaining reads using the tools built into GATK v4.0.8.1 (Van der Auwera et al., 2013). Subsequently, HaplotypeCaller and GentypeGVCFs were applied to call variants (SNPs and indels, respectively) per sample and used to perform joint genotyping. These results were used to generate a vcf file containing all raw SNPs and indels. SelectVariants and VariantFiltration tools in GATK were used to extract biallelic SNPs, which were subjected to hard-filtering for low-quality SNPs based on several scores (QD < 2.0, FS > 60.0, MQ < 40.0, MQRankSum < -12.5, ReadPosRankSum < -8.0, and SOR > 3.0). The distribution of each of the quality scores and their cut-offs was visualized in R (Supplemental Figure 11). Subsequently, SNPs were further filtered by Minor Allele Frequency (MAF) and the missing rate for each SNP (MAF < 0.05, missing rate > 0.1) for downstream analysis. Lastly, the filtered SNP call-set was annotated with SnpEff v4.3 using default settings to predict the nucleotide change effect of every SNP (Cingolani et al., 2012). Generated read data have been deposited in the European Nucleotide Achieve (ENA) under reference number PRJEB36060.

### Population structure analysis

PLINK2 was used to prune the SNP dataset to reduce the redundancy caused by linkage disequilibrium (LD) analysis for different downstream analyzes (Purcell et al., 2007). Firstly, SNPhylo was used to construct a maximum likelihood (ML) phylogenetic tree using 210,358 SNPs with default settings and 1,000 bootstrap replicates (window size = 50 SNPs, sliding size = 10, LD < 0.1) (Lee et al., 2014). Secondly, a PCA was conducted using PLINK2 on the pruned dataset of 904,930 SNPs (window size = 50 SNPs, sliding size = 10 SNPs, LD < 0.5). K-means clustering was performed via Eigen decomposition for the PCA and visualized in R. ADMIXTURE (v1.3.0) was used to deduce ancestral history and population structure using 96,804 SNPs (window size = 50 SNPs, sliding size = 10 SNPs, LD < 0.05) (Alexander and Lange, 2011). ADMIXTURE was utilized to determine the best number of ancestral populations (K = 1 to 4) by cross-validation errors (Supplemental Figure 12), and then was run again with the best K value with 1,000 bootstrap replicates to infer population structure. Population structure results were summarized using GISCO geographical information and visualized in R (Wickham, 2016; Eurostat GISCO, 2006).

### Comparative genomic analysis

The longest representative transcripts were selected from *L. saligna* and *L. sativa* as the basis for synteny analysis. BLAST (v2.2.31) was used to search homologous gene pairs between both species. MCScanX was employed to detect syntenic blocks (E-value cut-off = 1e-5, collinear block size ≥ 5) between the two *Lactuca* species using the top five alignment hits, which were visualized using SynVisio and JCVI (Tang et al., 2015a; Wang et al., 2012; Bandi and Gutwin, 2020). A separate synteny plot was created using the best-matching hit to remove noise from polyploidy and translocation events. Genetic markers on Chromosomes 5 and 8 were collected, and dot plots coordinated by genetic and physical positions were visualized with R v3.6.1 to validate inversions detected by synteny analysis. To assess the influence of genomic inversions, syntenic gene pairs located at inversion borders were searched against the *A. thaliana* protein database Araport11 (https://www.arabidopsis.org).

### NLR identification and classification

Genome-wide searches to identify *NLRs* were conducted using the genomes of *L. saligna* (v4) and *L. sativa* (v8) (downloaded from the CoGe website; gid35223). HMMER was used to search Hidden Markov Models (HMMs) for structural domains of NLRs (E-value cut-off = 1e-10). The Pfam models used were PF00931.23 and NBS_712.hmm for the NB domain, PF01582.20 and PF13676.6 for TIR, PF18052.1 for CC, and eight HMMs for the LRR domain (PF00560.33, PF07723.13, PF07725.13, PF12799.7, PF13306.6, PF13516.6, PF13855.6, PF14580.6). NB domains identified by InterProScan (see Functional annotation section) and CC motifs predicted by Paircoil2 (McDonnell et al., 2006) (P scores < 0.025) were integrated with the HMMER output. *NLRs* of *L. saligna* were classified into different categories (TNL/CNL and RGC families) by phylogeny clustering using *NLR*s previously identified in the *L. sativa* v8 genome (Christopoulou et al., 2015b). RGC families with a >3 count difference and 1.5 ratio between two species were selected as families with major differences. Amino acid sequences of NB domains were aligned with HmmerAlign (Finn et al., 2011). The alignment was trimmed by trimAl using the ’-gappyout’ algorithm, retaining 1,367 residues for phylogeny construction (Capella-Gutiérrez et al., 2009). The best-hit model of evolution, Blosum62+F+R10, was first selected by IQ-TREE v1.6.12 and ML trees were inferred with IQ-TREE (Nguyen et al., 2015). IQTREE (-pers 0.1, -nm 500) was run independently 10 times with 1,000 ultrafast bootstrap (UFBoot) replicates. Finally, the 10 best ML trees inferring the tree with highest log-likelihood was selected for NLR classification. Phylogenetic trees were visualized and annotated using iTOL v6 (Letunic and Bork, 2021).

### Identification of *NLR* clusters

Annotated *NLR* genes were used to determine gene intervals of MRCs on the *L. saligna* genome. The syntenic regions of MRCs in *L. saligna* were named lsal-MRCs to distinguish them from those detected in the *L. sativa* genome. An additional sliding window search was performed to identify *NLR* clusters (NCs) containing more than five *NLR*s (maximum10-genes gap). Identified MRCs and NCs were visualized on the *L. saligna* genome using Circos (Krzywinski et al., 2009).

### RLK identification and classification

Sequence similarity searches against primary protein sequences were performed with HMMER v3.1 using the PKinase alignment file (PF00069; E-value cut-off = 1e-10). Obtained protein sequences were subsequently scanned for the presence of extracellular domains using HMMER (E-value cut-off = 1e-3; Supplemental Table 23). TMHMM2.0 and SCAMPI2 were used to detect transmembrane regions (Krogh et al., 2001; Peters et al., 2016).

### Mapping NHR and HI regions

Genetic markers previously used in assembly reconstruction were aligned to genome assembly via BLAST v2.2.31 to locate the HI and NHRs regions in *L. saligna*. The genomic positions of one HI and four NHR regions were subsequently plotted on the *L. saligna* genome using Circos.

### RNA-seq analysis

Raw RNA-seq reads were quantified on *L. saligna* transcripts using Kallisto (v.0.44.0) to gain normalized transcript per million (TPM). Transcripts with a TPM value below 0.1 were considered not expressed. Then, DESeq2 was used to normalize the read count for each gene (total read count > 3) and execute statistical analyses to determine the DEGs with padj < 0.05 and log_2_FC > 1 (Love et al., 2014). Next, the read count mean and SD of infected and mock samples were calculated for all DEGs. Metascape was used for enrichment analysis of up-regulated genes and to render protein–protein interaction networks in Cytoscape (Zhou et al., 2019; Shannon et al., 2003). To identify potential candidate genes in the four NHR regions, additional counting for DE *RLK* and *NLR* genes, and highly regulated genes (|Log2FC| > 3) were counted separately.

## Data availability

The genome assembly described in this paper, *L. saligna* v4, is available under the BioProject PRJEB56287. All raw sequencing reads have been deposited in the ENA database under BioProject PRJEB56288. This includes the Illumina, PacBio, 10x Genomics, Bionano and Hi-C whole-genome sequences as well as RNA sequencing data for genome annotation and statistical analysis of *Bremia*-infection assay. The resequencing data for 15 *L. saligna* accessions are deposited under the BioProject PRJEB36060, which contains data of 100 *Lactuca* accessions derived from the TKI- 100 project.

## Author contributions

M.E.S, S.P, R.v.T, M.J conceived of the project; L.B, S.P, M.J, K.B and M.E.S designed experiments; F.RM.B, E.S, and LV.B generated plant material and sequencing data; L.B, and LV.B performed the genome assembling; W.X conducted the research and performed the analyses; W.X wrote the manuscript with assistance from other authors; all authors approved the final manuscript.

## ACKNOWLEDGMENTS

This research was supported by a grant of the International Lettuce Genomics Consortium (ILGC) funded by the Top Consortium for Knowledge and Innovation Horticultural and Starting Materials (grant number 1406-039). W.X. was financially supported by a fellowship by the China Scholarship Council (CSC). We thank Rens Holmer, Zhaodong Hao, Deedi Sogbohossou, Nora Walden, Sandra Smit, and Henri van de Geest for their support with bioinformatic analysis and genome sequencing data. We would also like to thank Wenhao Li for his help with statistics and R programming. We also thank Elizabeth Marie Georgian for writing support.

---

**Supplemental Figure 1.**
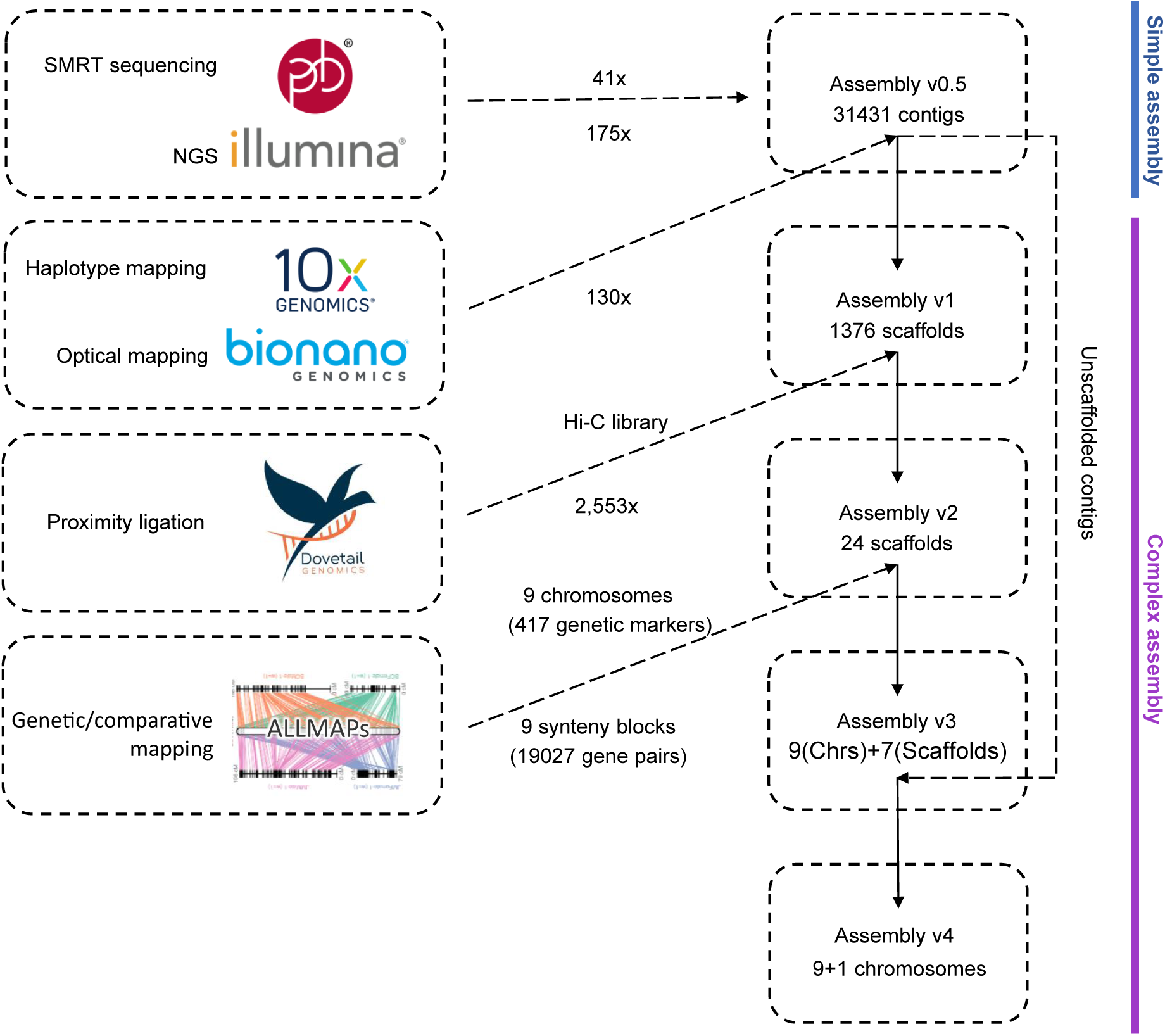
Sequencing and assembly workflow to construct the *L. saligna* reference genome. The sequencing techniques and reconstruction approaches are placed on the left. The dashed lines display contigs/scaffolds construction.

**Supplemental Figure 2.**
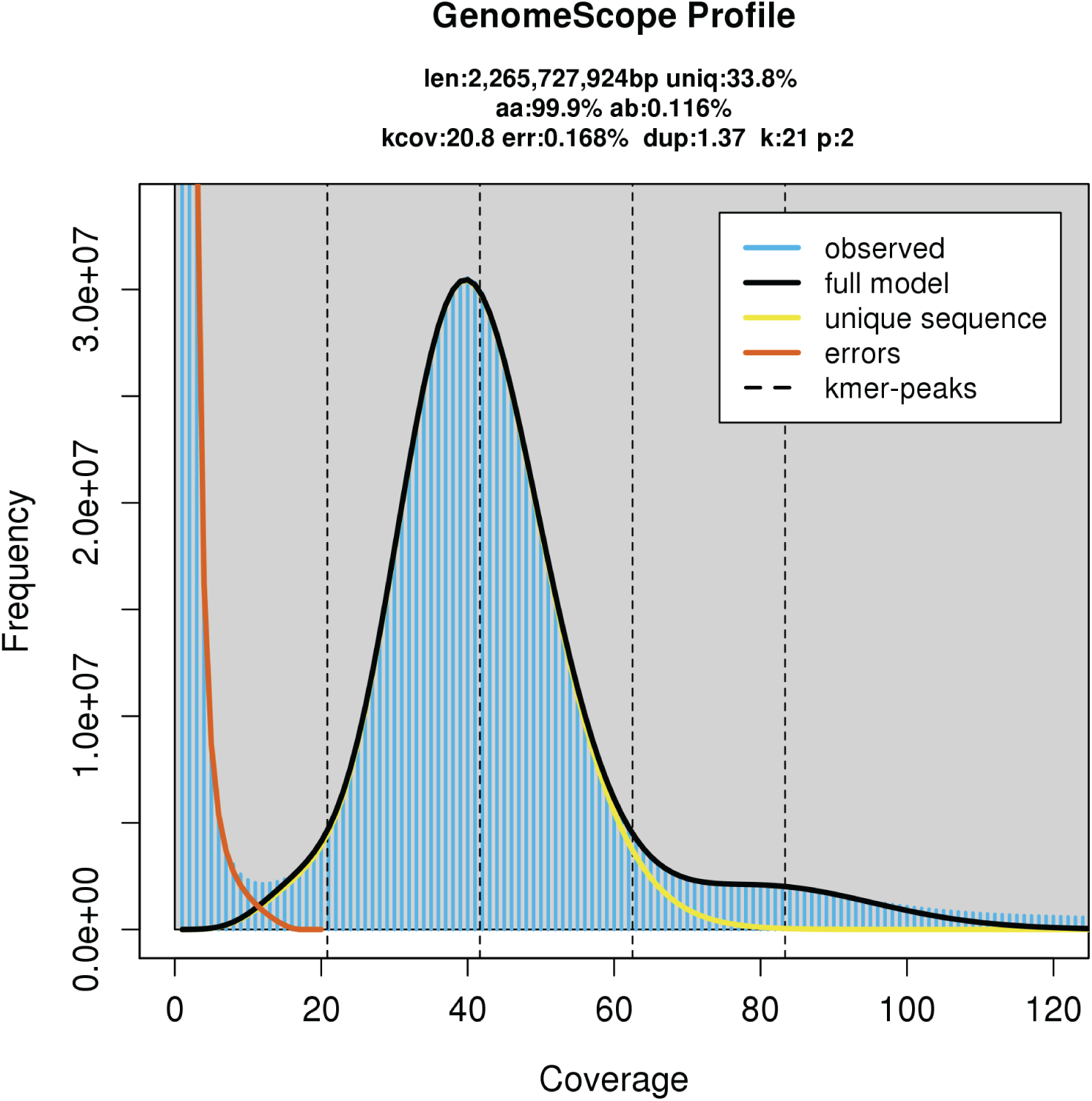
Genome size estimation of *L. saligna* by GenomeScope. The 21-mers were counted by Jellyfish, and output was taken by GenomeScope to estimate the genome size of *L. saligna*. The frequency (y-axis) and sequencing depth (x-axis) of 21-mer are plotted. The genome size (2.27 Gb) was estimated by the highest peak depth. len: Genome haploid length; uniq: genome uniq length; aa: homozygosity %; kcov: k-mer coverage; err: read error rate; dup: average rate of read duplication; k: k-mer length; p: ploidy.

**Supplemental Figure 3.**
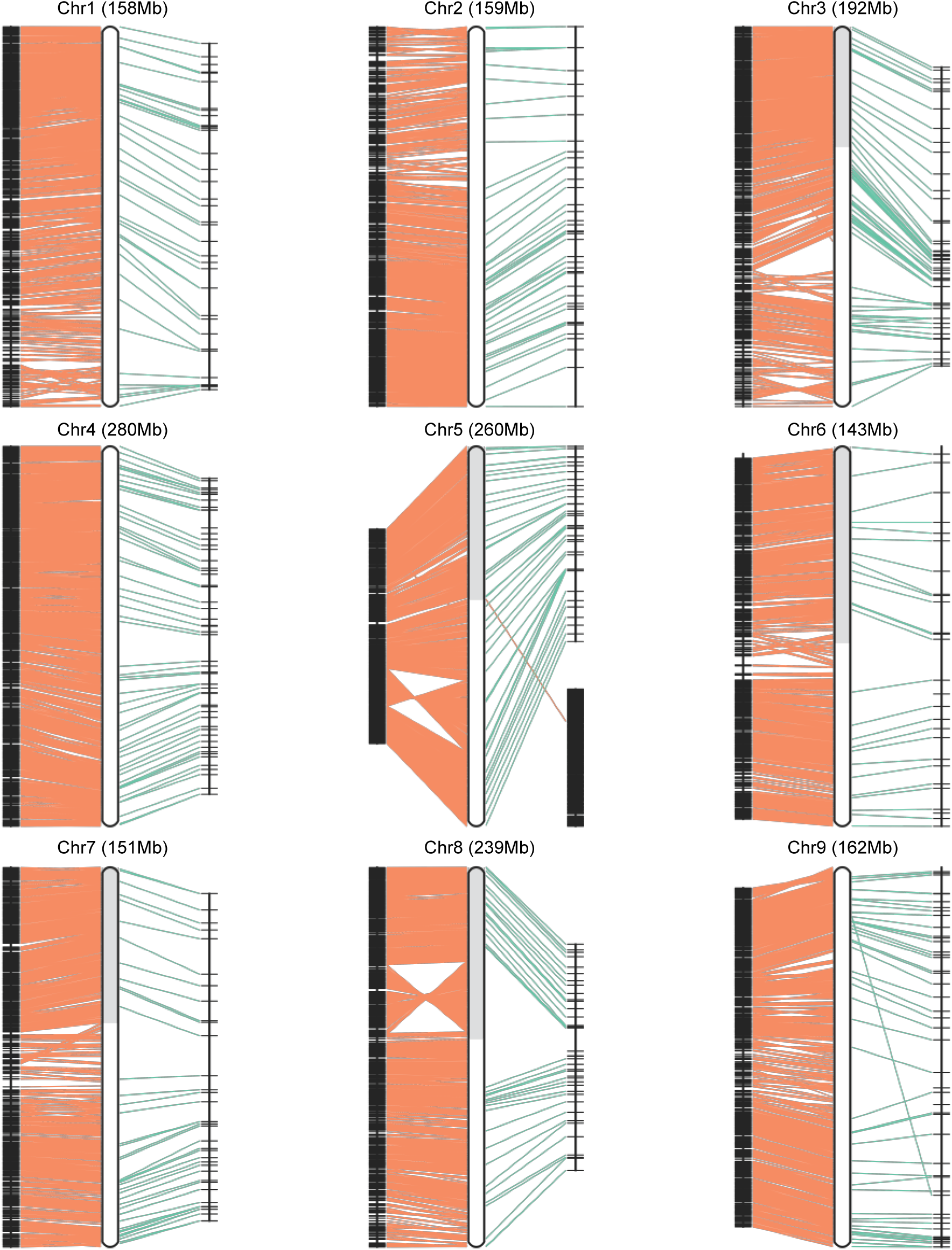
ALLMAPS re-scaffolding for *L. saligna* pseudo-chromosomes using genetic and syntenic map. The *L. saligna* assembly v3 was reconstructed into 9 pseudo-chromosomes using genetic markers (weight=2) and syntenic markers (weight=1). For linkage map of each chromosome, left bar represents genes from the same synteny block, the right bar indicates the markers’ position on genetic map. The two types of markers are connected to the middle ideogram of each reconstructed chromosome by orange and green lines respectively. Grey and white colors in chromosomes stand for different scaffolds from previous assembly.

**Supplemental Figure 4.**
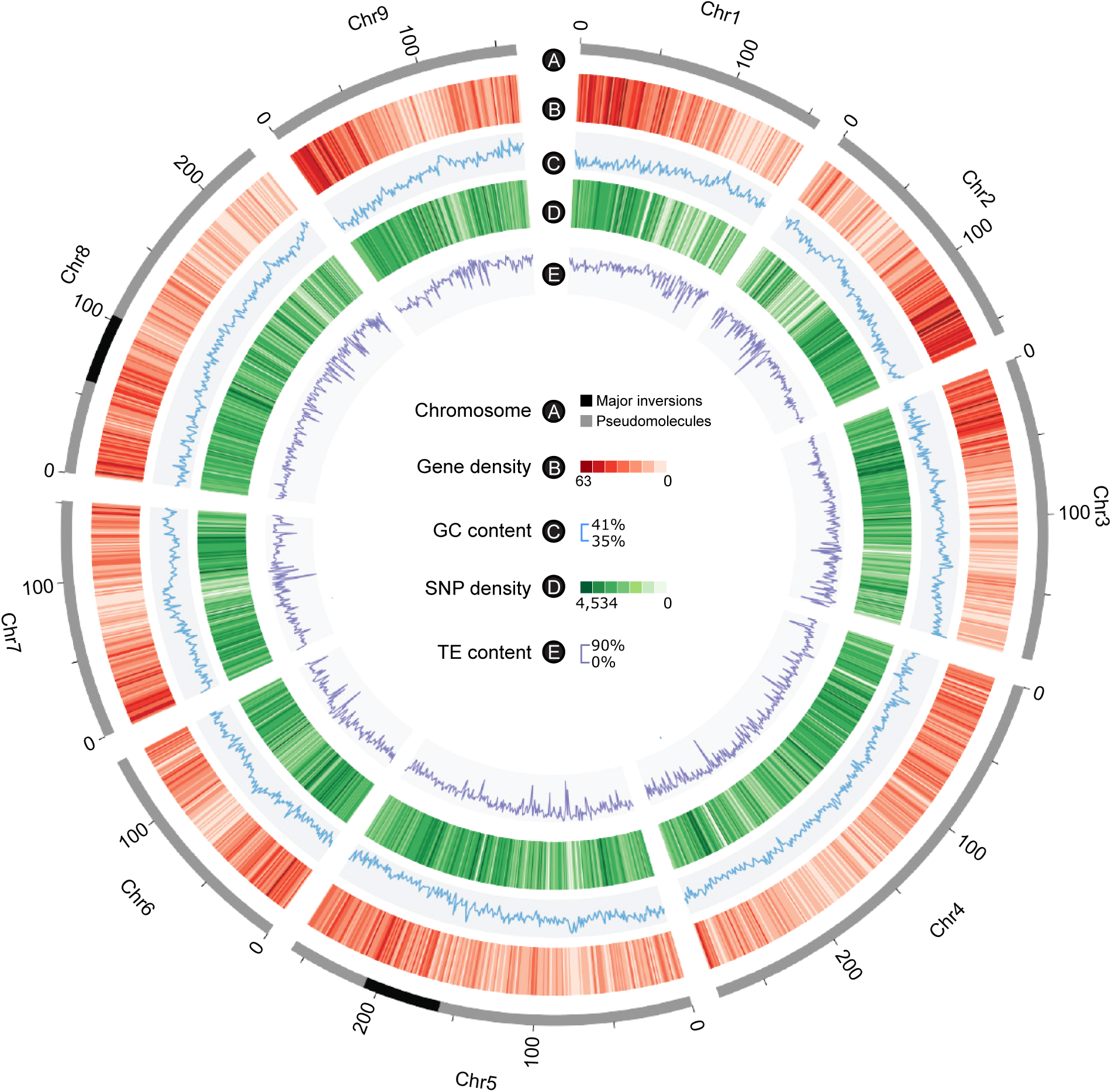
Genomic features of the *Lactuca saligna* genome. A, Circular ideogram of the nine pseudo-chromosomes of *L. saligna* CGN05327 at Mb scale. Grey areas represent oriented regions compared to *L. sativa*. B, black regions represent inverted regions between *L. saligna* and *L. sativa*. B, Gene density (red; 1Mb window). C, GC content percentage (blue line; 1Mb window; outward). D, Density of single-nucleotide polymorphisms (SNPs) (green; 1Mb window). E, TE content percentage (purple line; 1Mb window; outward).

**Supplemental Figure 5.**
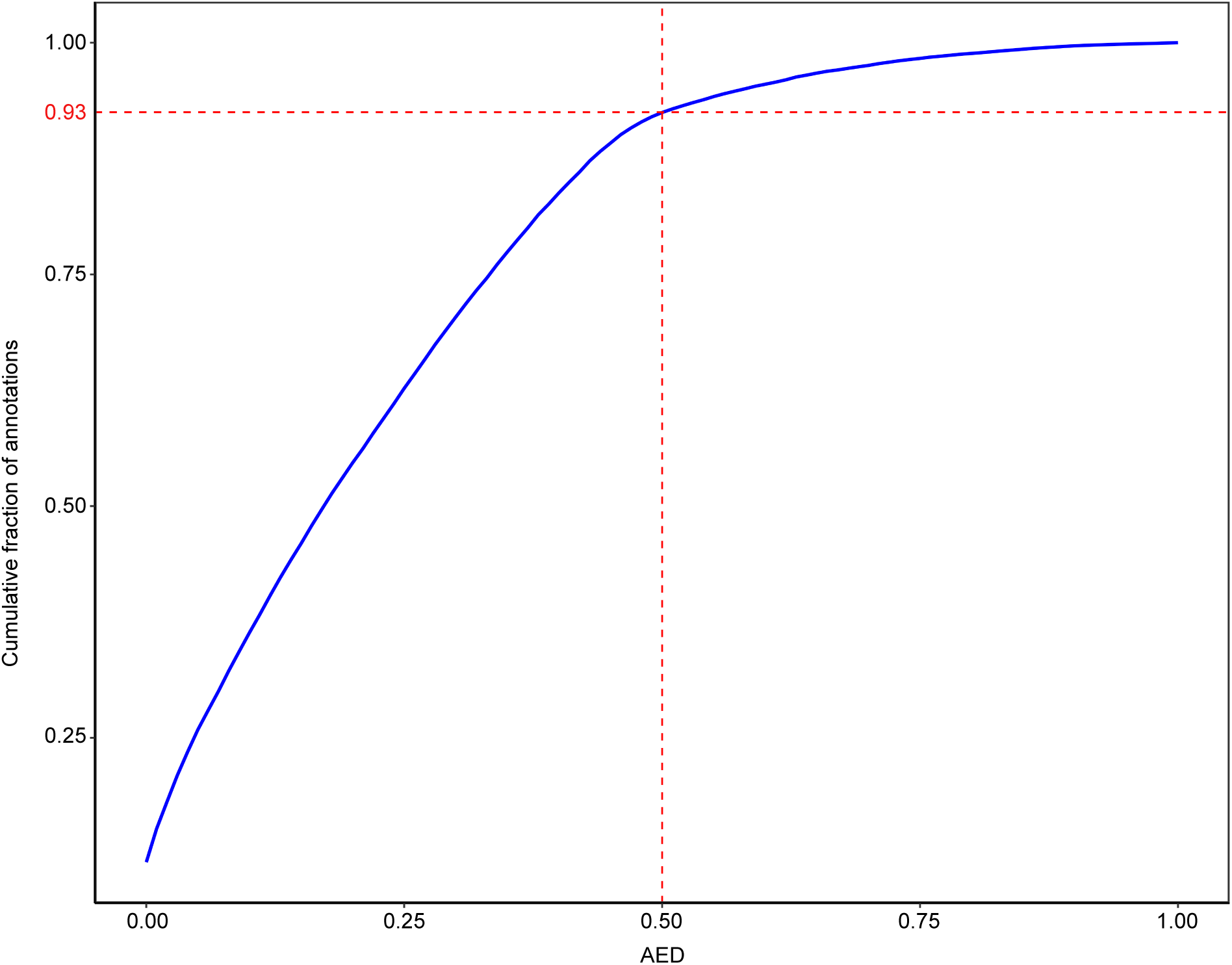
Annotation Edit Distance (AED) cumulative fraction genome *de novo* annotation. Annotation Edit Distance (AED) indicates how well a predicted gene model is supported by biological evidence. AED values range from 0 and 1, with 0 denoting perfect agreement of the annotation to aligned evidence, and 1 denoting no evidence support for the annotation. Around 93% of the annotations having AEDs of less than 0.5, where AED smaller than 0.5 indicates a well-supported gene model.

**Supplemental Figure 6.**
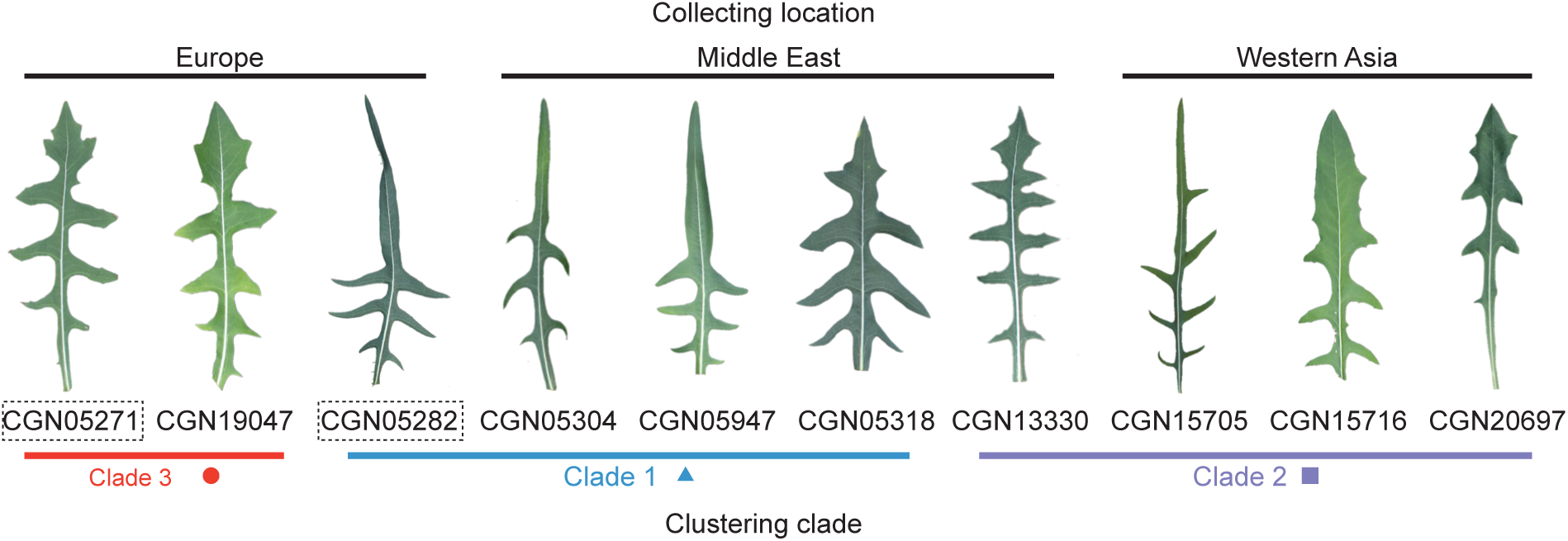
Diversity of leaf shape of 10 resequenced *L. saligna* accessions. Leaf shapes of *L. saligna* accessions of European, Middle Eastern and West Asian origin reordered by clustering result (Figure 1).

**Supplemental Figure 7.**
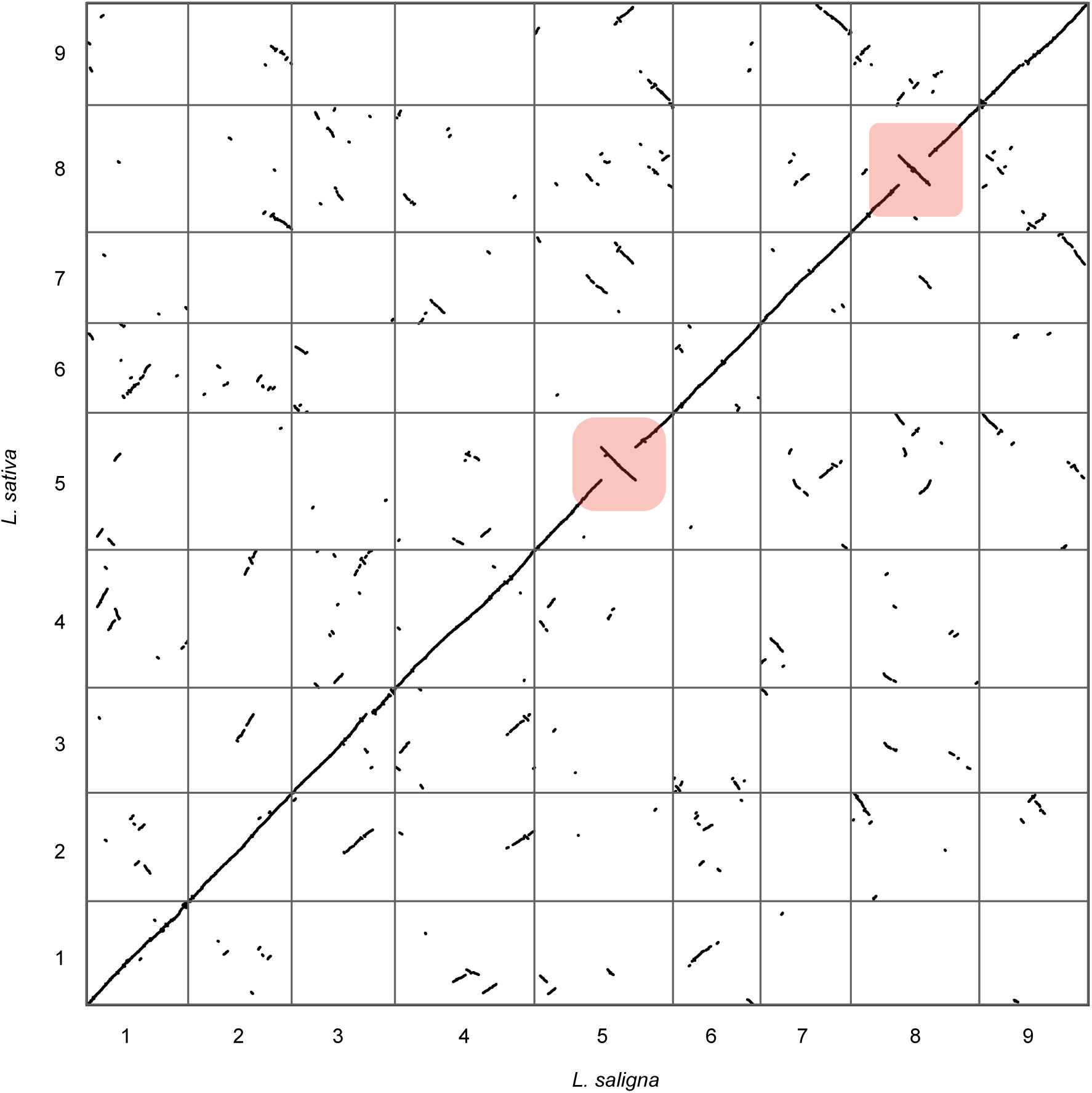
Syntenic path dot plot of *L. sativa* versus *L. saligna* highlighting two large inversions on chromosomes 5 and 8. The y-axis represents the 9 *L. sativa* chromosomes, the x-axis represents the 9 *L. saligna* chromosomes. Overall, primary synteny is seen between all chromosomes (major diagonal line). Some off-axis synteny is also seen due to the ancient shared polyploidy between these two species. The inverted synteny blocks on chromosomes 5 and 8 are highlighted by pink boxes.

**Supplemental Figure 8.**
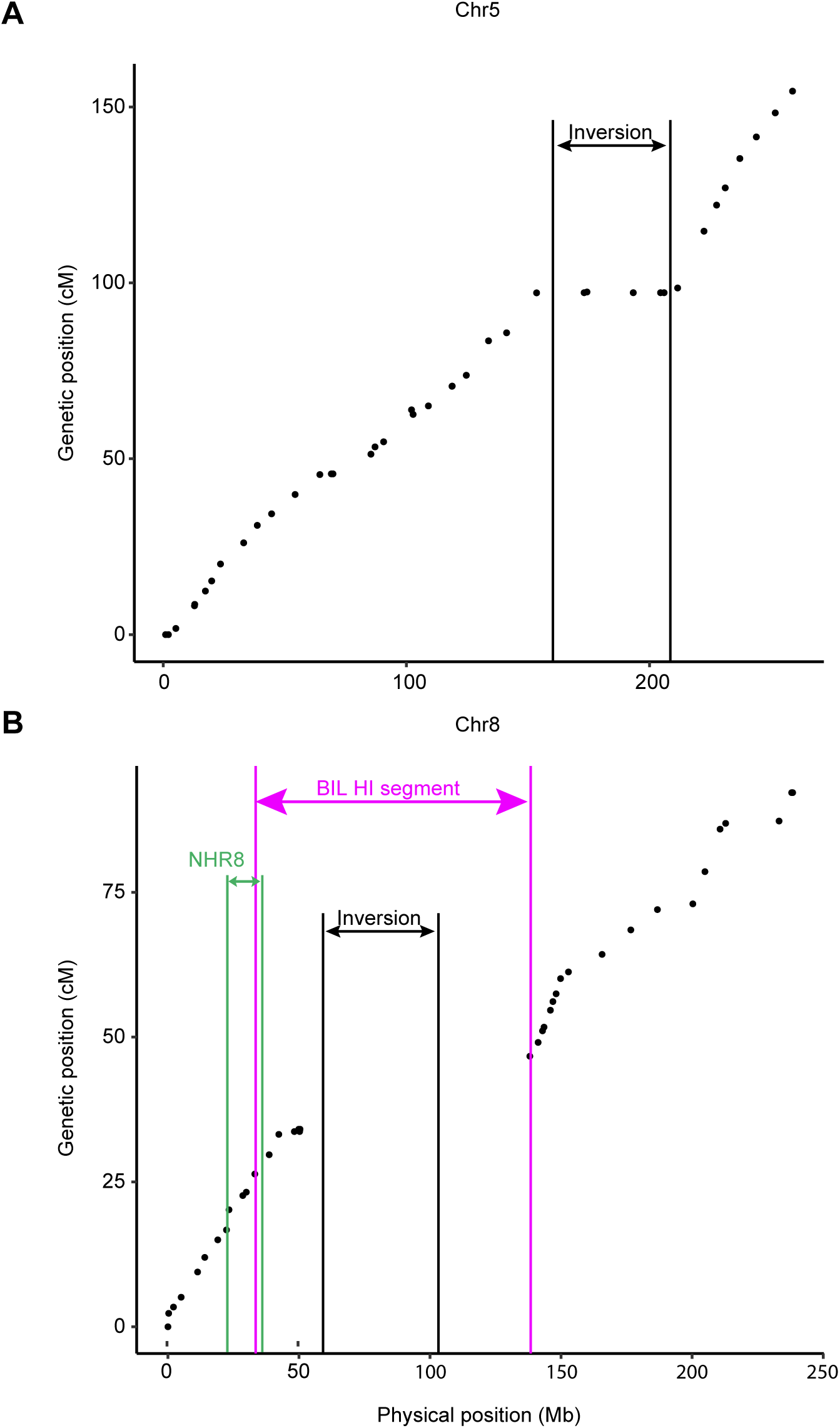
Genetic distance versus physical position on chromosome 5 and 8 supporting the presence of large inversions (i.e. a small genetic distance corresponding to a large physical distance). The SNP-derived markers for chromosome reconstruction were used for plotting genetic distance (based on an F2 genetic map of *L. saligna* x *L. sativa*) versus physical position in *L. saligna* on chromosome 5 (A) and 8 (B). The y-axis represents the genetic position of the genetic markers. The x-axis represents the physical position of genetic markers on the *L. saligna* genome. The inversion intervals are denoted by black lines based on the physical position of syntenic genes. The Hybrid Incompatibility (HI) regions found by TRD in BIL is located by pink line. The locus of Nonhost Resistance (NHR) fine-mapped on chromosome 8 (NHR8) is denoted by green line (Support Figure 3).

**Supplemental Figure 9.**
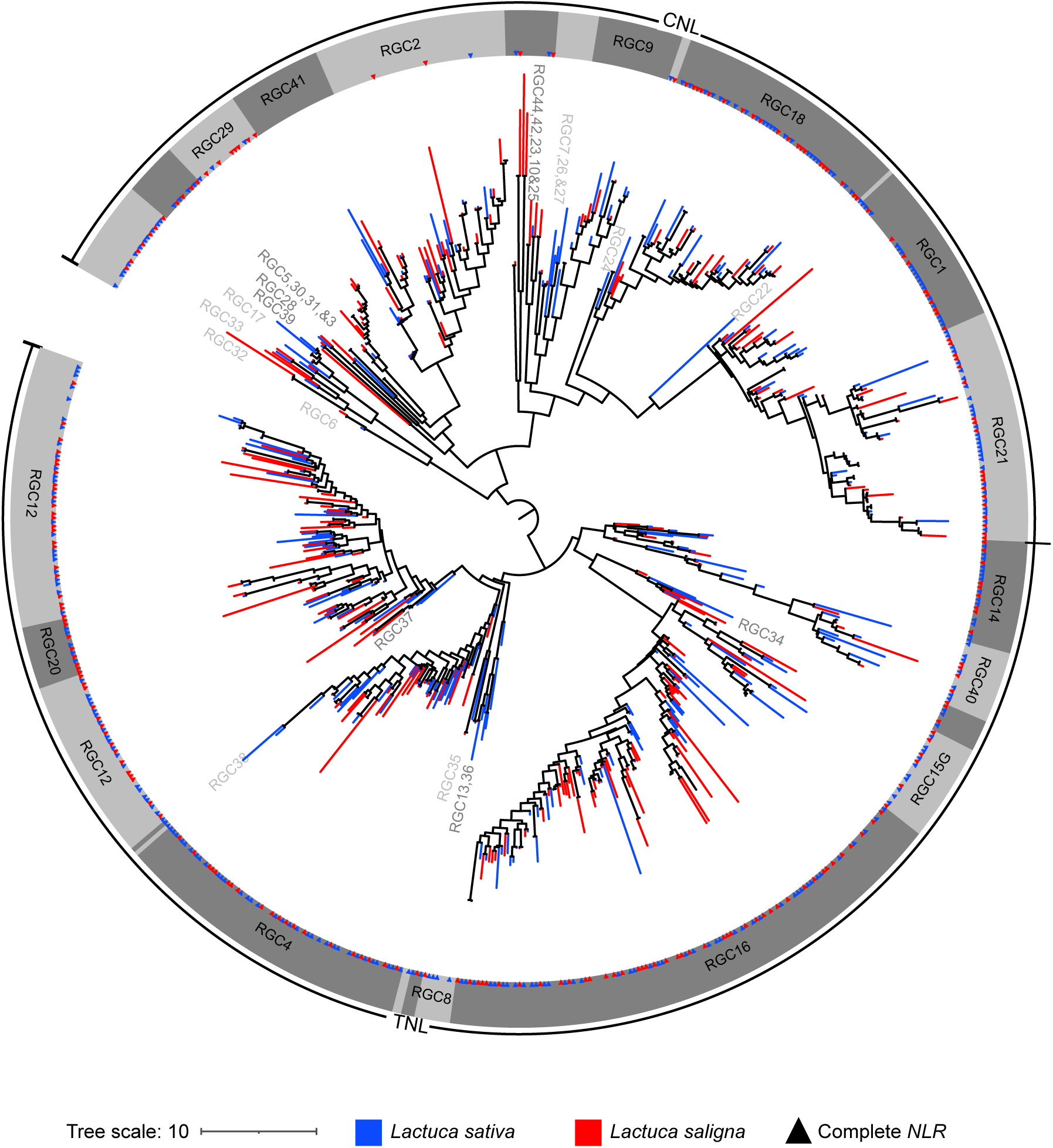
Circular tree of *L .saligna* nucleotide binding-leucine rich repeat receptors (NLR) generated by IQTREE. The tree was re-rooted at the midpoint between TNL and CNL clade, and ultra- fast bootstrap approximation (UFBoot) support values was calculated 1000 repetitions. Branch color represents the species: blue for *L. sativa*, read for *L. saligna*. Triangle indicates the *NLR*s with complete domain structure via HMMER search. Nomenclature of resistance gene candidate (RGC) family was referring to previously phylogeny of lettuce (Christopoulou et al., 2015b).

**Supplemental Figure 10.**
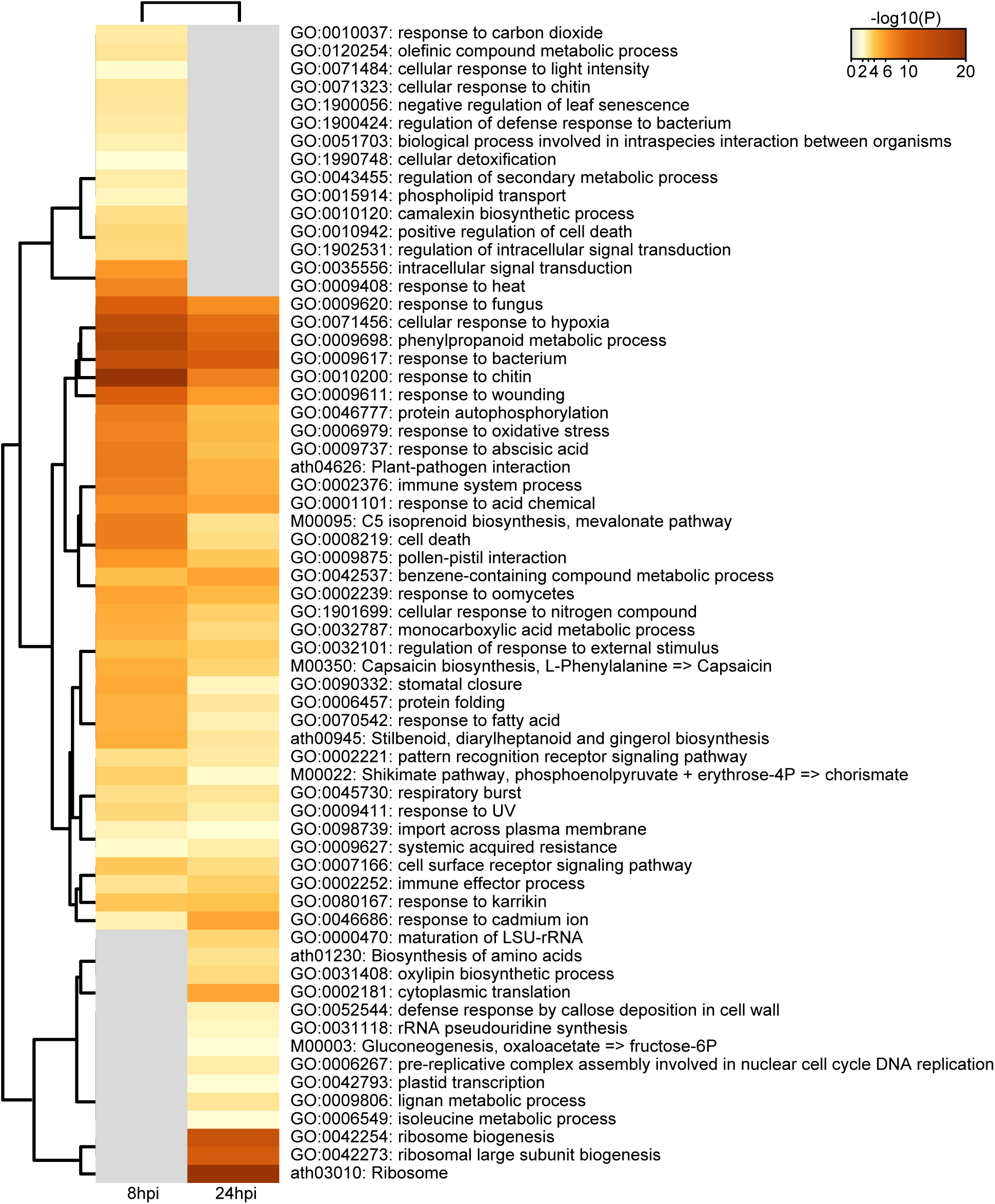
Top 100 enriched biological clusters of up-regulated DE genes at 8 and 24 hpi. The enrichment analysis of ontology clustered the up-regulated DE genes at 8 and 24 hpi. The *Arabidopsis thaliana* gene ids were used to annotate DE genes and the plots were adjusted from the enrichment analysis results generated by Metascape platform. Heatmap of the top 100 ontology groups represented by the term with the best p-value. The terms are ordered by the hierarchically clustering result. The heatmap cells are colored by transformed p-values. Grey cells indicate the lack of enrichment for that term in the corresponding gene list. The number of each cluster or row in heatmap represents the order of clustering result.

**Supplemental Figure 11.**
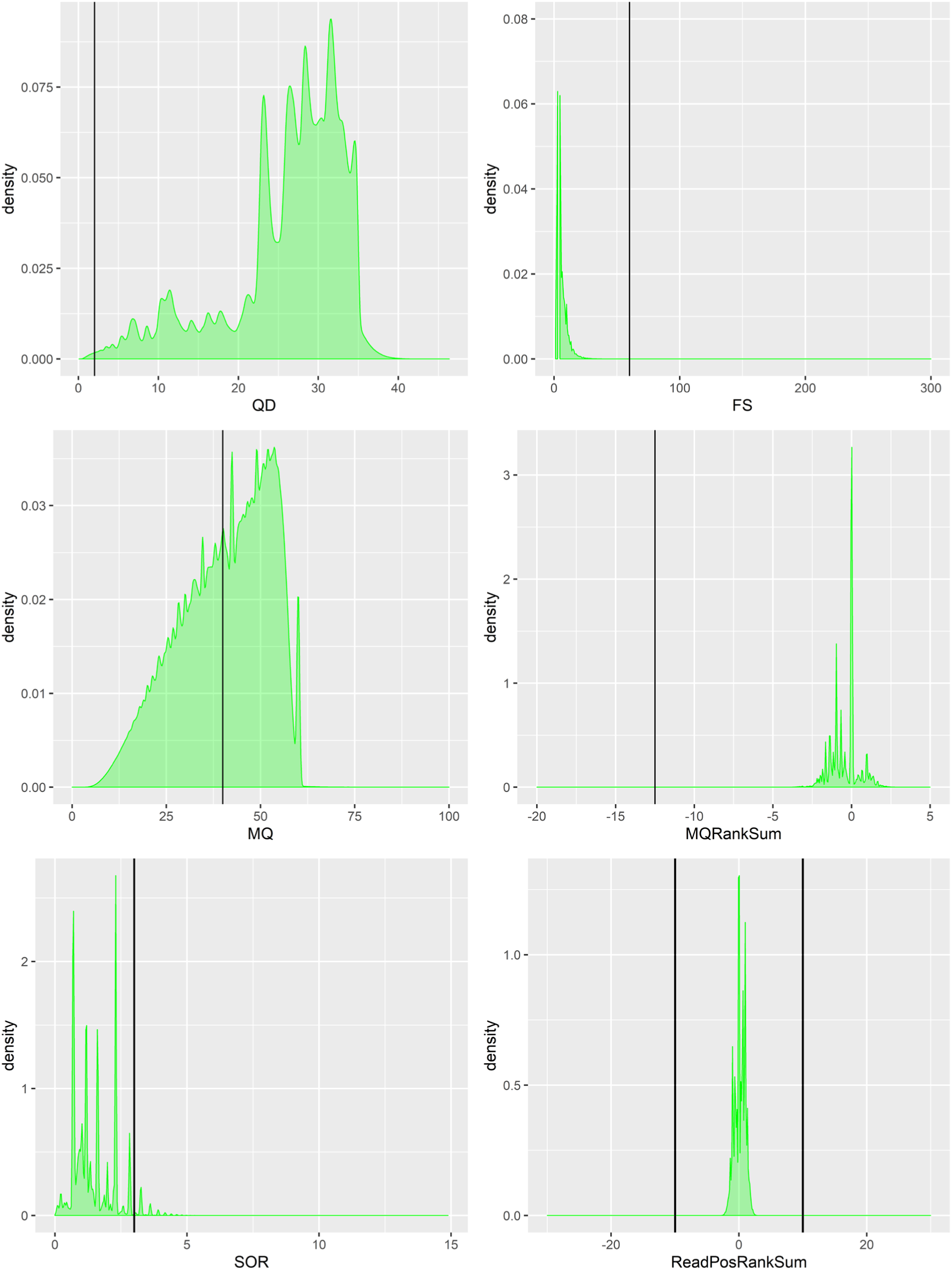
Hard-filtering for biallelic SNPs variant from resequencing analysis of 15 *L. saligna* accessions. The distribution of six quality parameters: QualByDepth (QD), FisherStrand (FS), RMSMappingQuality (MQ), MappingQualityRankSumTest (sMQRankSum), StrandOddsRatio (SOR), ReadPosRankSumTest (ReadPosRankSum). The filtering cut-off values are denoted by black lines.

**Supplemental Figure 12.**
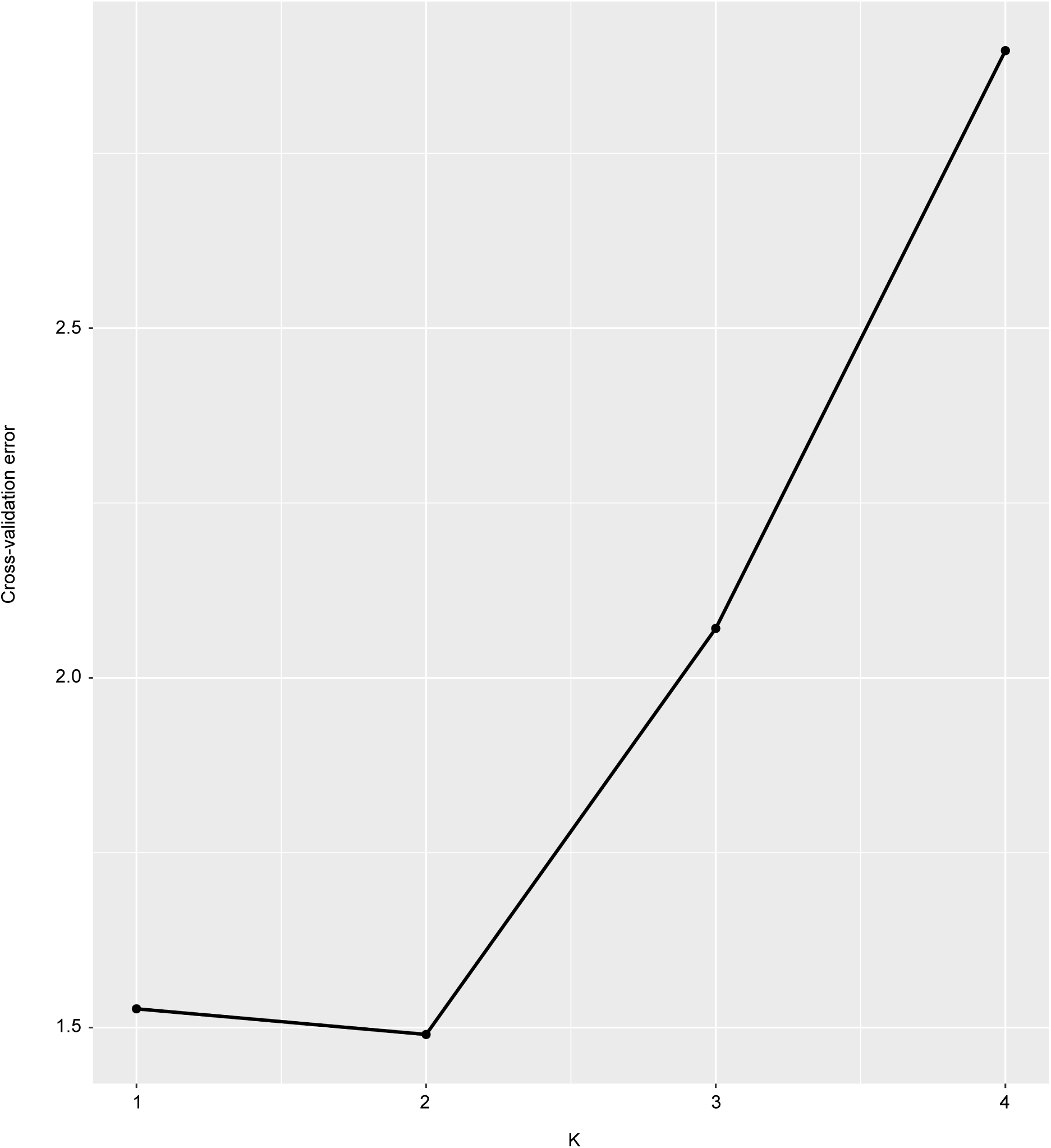
Cross-validation estimates the best K (population numbers) of ADMIXTURE for 15 re-sequenced *L. saligna* accessions. A good number of population (K) has a lower cross-validation (CV) error compared to other K values. K value ranges from 1 to 4 were applied to calculate the CV error. The K with lowest CV error was assumed as the best estimation for number of ancestral populations.

## SUPPLEMENTAL TABLES

**Supplemental Table 1** Resequencing data for genome size estimation

**Supplemental Table 2** ALLMAPS genome reconstruction by genetic and syntenic mapping

**Supplemental Table 3** ALLMAPS summary for the consensus map

**Supplemental Table 4** Chromosomal length of *L. saligna* (v4)

**Supplemental Table 5** BUSCO assessment*

**Supplemental Table 6** Mapping ESTs of different *Latuca* species to *L. saligna* genome

**Supplemental Table 7** Repeat annotation summary

**Supplemental Table 8** Categories of TEs predicted in the *L. saligna* genome

**Supplemental Table 9** Subcategories of TEs predicted in the *L. saligna* genome

**Supplemental Table 10** ncRNA prediction by different tools

**Supplemental Table 11** Functional gene annotation

**Supplemental Table 12** Passport of 15 *L. saligna* accessions

**Supplemental Table 13** Summary of *L. saligna* re-sequencing

**Supplemental Table 14** Summary of filtered SNPs for 15 TKI lines against *L. saligna* reference genome (v4)

**Supplemental Table 15** Annotation of filtered SNPs by SnpEff

**Supplemental Table 16** Chromosomal position and gene count of inversions

**Supplemental Table 17** Genes at the border of putative breaking regions

**Supplemental Table 18** Genes flanking interspecific inversions between *L. saligna* and *L. sativa*

**Supplemental Table 19** ID conversion of genes flanking interspecific inversion

**Supplemental Table 20** *NLR* gene number in *L. saligna* and *L. sativa*

**Supplemental Table 21** Genomic position of Major Resistance Clusters (MRCs) in *Lactuca saligna*

**Supplemental Table 22** *NLR* clusters (NCs) in the *Lactuca saligna* v4 assembly

**Supplemental Table 23** Number of *NLR*s per RGC family in *L. sativa* and *L. saligna*

**Supplemental Table 24** Pfam HMM motifs used for *RLK* classification

**Supplemental Table 25** Hybrid incompatibility (HI) regions mapping in *L. saligna v4 assembly*

**Supplemental Table 26** NHR intervals and *R* gene locus mapping in *L. saligna* v4 assembly

**Supplemental Table 27** Number of differentially expressed genes in *L. saligna* upon *Bremia* infection

**Supplemental Table 28** DEGs in oomycete related ontology of Bremia-infected *L. saligna*

**Supplemental Table 29** Expression level of DEGs in oomycete related ontology of Bremia-infected *L. saligna*

**Supplemental Table 30** Statistics of DEGs in four NHR regions of *L. saligna*

**Supplemental Table 31** Candidate genes in NHR8 with potential *L. saligna* nonhost resistance

## SUPPLEMENTAL DATASETS

Supplemental Dataset 1

Supplemental Datasets 2A-F

Supplemental Datasets 3A-B

Supplemental Datasets 4A-B

Supplemental Datasets 5A-D

Supplemental Datasets 6A-B

Supplemental Dataset 7

