## Supplemental Tables for "The genome of *Lactuca saligna*, a wild relative of lettuce, provides insight into non-host resistance to the downy mildew *Bremia lactucae*"

**Supplemental Table 1** Resequencing data for genome size estimation

| **fastq** | ***n* of reads** | **total length (bp)** |
| --- | --- | --- |
| run0161IL_*Lactuca*_saligna_500_S16_L007_R1_001.nophix.fastq | 232,811,628 | 29,212,084,397 |
| run0161IL_*Lactuca*_saligna_500_S16_L007_R2_001.nophix.fastq | 232,811,628 | 29,212,174,566 |
| run0161IL_*Lactuca*_saligna_500_S16_L008_R1_001.nophix.fastq | 233,145,034 | 29,253,891,835 |
| run0161IL_*Lactuca*_saligna_500_S16_L008_R2_001.nophix.fastq | 233,145,034 | 29,253,857,540 |
| Total | 931,913,324 | 116,932,008,338 |

**Supplemental Table 2** ALLMAPS genome reconstruction by genetic and syntenic mapping

| **Metrics** | **Genetic marker** | **Syntenic marker** |
| --- | --- | --- |
| Linkage Groups | 9 | 9 |
| Markers (unique) | 417 | 19,027 |
| Markers per Mb | 0.2 | 10.9 |
| N50 Scaffolds | 5 | 5 |
| Scaffolds | 16 | 17 |
| Scaffolds with 1 marker | 1 | 0 |
| Scaffolds with 2 markers | 0 | 0 |
| Scaffolds with 3 markers | 0 | 0 |
| Scaffolds with >=4 markers | 15 | 17 |
| Total bases | 1,728,492,023 (98.9%) | 1,745,017,402 (99.8%) |

**Supplemental Table 3** ALLMAPS summary for the consensus map

|  | **Anchored** | **Oriented** | **Unplaced** |
| --- | --- | --- | --- |
| Markers (unique) | 19,443 | 19,443 | 0 |
| Markers per Mb | 11.1 | 11.1 | 0 |
| N50 Scaffolds | 5 | 5 | 0 |
| Scaffolds | 17 | 17 | 7 |
| Scaffolds with 1 marker | 0 | 0 | 0 |
| Scaffolds with 2 markers | 0 | 0 | 0 |
| Scaffolds with 3 markers | 0 | 0 | 0 |
| Scaffolds with >=4 markers | 17 | 17 | 0 |
| Total bases | 1,745,017,402 (99.8%) | 1,745,017,402 (99.8%) | 3,335,274 (0.20%) |

**Supplemental Table 4** Chromosomal length of *L. saligna* (v4)

| **Chromosome** | **Length (bp)** |
| --- | --- |
| chr1 | 158,169,978 |
| chr2 | 159,493,583 |
| chr3 | 192,140,685 |
| chr4 | 279,860,179 |
| chr5 | 259,520,967 |
| chr6 | 143,320,011 |
| chr7 | 151,383,248 |
| chr8 | 238,633,233 |
| chr9 | 162,496,318 |
| chr0 | 420,743,833 |

**Supplemental Table 5** BUSCO assessment*

| **BUSCO categories** | **Count** | **%** |
| --- | --- | --- |
| Complete BUSCOs | 1,951 | 91.9 |
| Complete and single-copy BUSCOs | 1,859 | 87.6 |
| Complete and duplicated BUSCOs | 92 | 4.3 |
| Fragmented BUSCOs | 44 | 2.1 |
| Missing BUSCOs | 126 | 6.0 |

**Supplemental Table 6** Mapping ESTs of different *Lactuca* species to *L. saligna* genome

| **Species** | ***n* of EST** | **80% identity** | | | | **50% identity** | | | |
| --- | --- | --- | --- | --- | --- | --- | --- | --- | --- |
|  |  | **80% coverage** | | **50% coverage** | | **80% coverage** | | **50% coverage** | |
|  |  | ***n*** | **%** | ***n*** | **%** | ***n*** | **%** | ***n*** | **%** |
| *Lactuca* spp. | 226,897 | 199,027 | 87.7 | 200,040 | 88.2 | 213,004 | 93.9 | 214,741 | 94.6 |
| *L. sativa* | 81,518 | 72,006 | 88.3 | 72,324 | 88.7 | 77,095 | 94.6 | 77,645 | 95.2 |
| *L. serriola* | 55,490 | 49,504 | 89.2 | 49,835 | 89.8 | 52,823 | 95.2 | 53,334 | 96.1 |
| *L. saligna* | 30,696 | 28,243 | 92.0 | 28,286 | 92.1 | 29,371 | 95.7 | 29,476 | 96.0 |
| *L. virosa* | 30,068 | 25,898 | 86.1 | 26,008 | 86.5 | 27,772 | 92.4 | 27,971 | 93.0 |
| *L. perennis* | 29,125 | 23,376 | 80.3 | 23,587 | 81.0 | 25,943 | 89.1 | 26,315 | 90.4 |

* Collected from NCBI (retrieved at 2019-7-11).

**Supplemental Table 7** Repeat annotation summary

| **Repeat type** | **Search type** | **Repeat Size (bp)** | **% of genome** |
| --- | --- | --- | --- |
| Transposable elements (TEs) | Homolog^a^ | 56,775,417 | 2.62 |
|  | Identified *de novo*^b^ | 1,536,393,161 | 70.94 |
|  | Unknown *de novo*^b^ | 143,637,521 | 6.63 |
|  | Merge^c^ | 1,678,347,818 | 77.49 |
| Tandem repeats^d^ |  | 72,446,575 | 3.35 |
| Total |  |  | 80.84 |

^a^ By RepeatMasker using Repbase and Dfam (Bao, Kojima and Kohany, 2015; Hubley *et al.*, 2015; Smit, Hubley and Green, 2019).

^b^ By RepeatMasker using the *de novo* transposon prediction (Price, Jones and Pevzner, 2005; Han and Wessler, 2010; Smit, Hubley and Green, 2019).

^c^ By combining serially identified repeats and removing redundancy.

^d^ Prediction using TRF (Benson, 1999).

**Supplemental Table 8** Categories of TEs predicted in the *L. saligna* genome

|  | **Repbase + Dfam** | |  | **Classified *de novo*** | |  | **Unknown *de novo*** | |  | **Combined TEs** | |
| --- | --- | --- | --- | --- | --- | --- | --- | --- | --- | --- | --- |
|  | **Length (Mb)** | **% in genome** |  | **Length (Mb)** | **% in genome** |  | **Length (Mb)** | **% in genome** |  | **Length (Mb)** | **% in genome** |
| DNA | 0.24 | 0.01 |  | 69.97 | 3.23 |  | 0 | 0 |  | 66.90 | 3.09 |
| LINE | 1.75 | 0.08 |  | 5.17 | 0.24 |  | 0 | 0 |  | 6.89 | 0.32 |
| SINE | 0.00 | 0.00 |  | 0.27 | 0.01 |  | 0 | 0 |  | 0.27 | 0.01 |
| LTR | 54.79 | 2.53 |  | 1,460.97 | 67.46 |  | 0 | 0 |  | 1,460.65 | 67.44 |
| Unknown | 0.00 | 0.00 |  | 0.00 | 0.00 |  | 143.64 | 6.63 |  | 143.64 | 6.63 |
| Total^a^ | 56.78 | 2.62 |  | 1,536.39 | 70.94 |  | 143.64 | 6.63 |  | 1,678.35 | 77.49 |

a. Different libraries were fed to RepeatMasker to search transposable elements in a serial order. There are some overlaps between different searches, the combined number is therefore less than the metric sum.

**Supplemental Table 9** Subcategories of TEs predicted in the *L. saligna* genome

| **Type** | **Subcategory** | **Copy-number** | **Length (bp)** | **% in genome** |
| --- | --- | --- | --- | --- |
| LTR | Total | 1,196,201 | 1,460,649,948 | 67.44 |
|  | Cassandra | 2,154 | 715,047 | 0.03 |
|  | Caulimovirus | 988 | 1,674,216 | 0.08 |
|  | Copia | 351,651 | 500,114,293 | 23.09 |
|  | Gypsy | 827,409 | 947,670,181 | 43.76 |
|  | Pao | 3,741 | 1,681,181 | 0.08 |
|  | Other | 10,258 | 8,795,030 | 0.41 |
| DNA | Total | 201,125 | 66,895,899 | 3.09 |
|  | CMC-EnSpm | 9,705 | 5,148,307 | 0.24 |
|  | Crypton-S | 572 | 142,627 | 0.01 |
|  | hAT-Ac | 14,294 | 4,565,181 | 0.21 |
|  | hAT-Charlie | 756 | 232,662 | 0.01 |
|  | hAT-Tag1 | 5,972 | 1,735,044 | 0.08 |
|  | hAT-Tip100 | 5,312 | 2,508,379 | 0.12 |
|  | MITE | 120,287 | 29,161,599 | 1.35 |
|  | MULE-MuDR | 19,874 | 12,616,217 | 0.58 |
|  | PIF-Harbinger | 11,418 | 6,269,922 | 0.29 |
|  | Helitron | 7,480 | 2,492,434 | 0.12 |
|  | TcMar-Stowaway | 518 | 95,638 | 0.00 |
|  | Other | 4,937 | 1,927,889 | 0.09 |
| LINE | Total | 10,015 | 6,890,512 | 0.32 |
|  | CR1 | 565 | 598,086 | 0.03 |
|  | CRE | 2,431 | 1,770,999 | 0.08 |
|  | CRE-II | 35 | 3,753 | 0.00 |
|  | L1 | 5,888 | 4,241,171 | 0.20 |
|  | RTE-BovB | 1,096 | 276,503 | 0.01 |
| SINE | Total | 1,686 | 273,938 | 0.01 |
|  | tRNA | 1,686 | 273,938 | 0.01 |
| Unknown | Total | 588,393 | 143,637,521 | 6.63 |

**Supplemental Table 10** ncRNA prediction by different tools

|  | **Specific** | **General^c^** | **Reduced^d^** |
| --- | --- | --- | --- |
| tRNA | 1,797^a^ | 1,793 | 1,857 |
| rRNA | 330^b^ | 4,168 | 4,114 |
| miRNA | - | 128 | 128 |
| snRNA | - | 329 | 329 |

^a^ Predicted by tRNAscan-SE (Lowe and Eddy, 1997).

^b^ Predicted by RNAmmers (Lagesen *et al.*, 2007).

^c^ By INFERNAL searching all types of ncRNA in Rfam database (Nawrocki and Eddy, 2013; Kalvari *et al.*, 2018).

^d^ Combination reduced by GenomicRanges (Lawrence *et al.*, 2013).

**Supplemental Table 11** Functional gene annotation

|  | ***n*** | **%** |
| --- | --- | --- |
| Total genes | 42,908 | 100.0 |
| Total annotated | 40,730 | 94.9 |
| InterPro | 27,373 | 63.8 |
| SwissProt | 27,327 | 63.7 |
| TrEMBL | 40,622 | 94.7 |
| KEGG | 13,426 | 31.3 |
| TAIR | 31,472 | 73.3 |
| Non-matched | 2,178 | 5.1 |

**Supplemental Table 12** Passport of 15 *L. saligna* accessions

| **Accession** | **SSD** | **latitude** | **longitude** | **Origin/Collection** | **Region** |
| --- | --- | --- | --- | --- | --- |
| CGN05271^a a^ | TKI-342 | 47.2 | -1.5 | JBN^b^ | Europe |
| CGN05282^a^ | TKI-343 | 47.2 | -1.5 | JBN^b^ | Europe |
| CGN05301 | TKI-344 | 44.5 | 1.5 | France | Europe |
| CGN05304^a^ | TKI-345 | 32.9 | 35.1 | Israel | Middle East |
| CGN05318^a^ | TKI-355 | 33.0 | 35.3 | Israel | Middle East |
| CGN05327 | TKI-364 | 41.9^c^ | 2.8^c^ | Spain | Europe |
| CGN05330 | TKI-366 | 31.5 | 35.1 | Israel | Middle East |
| CGN05947^a^ | TKI-369 | 33.0 | 35.5 | Israel | Middle East |
| CGN13326 | TKI-374 | 38.0 | 23.9 | Greece | Europe |
| CGN13330^a^ | TKI-376 | 37.4 | 27.6 | Turkey | West Asia |
| CGN13375 | TKI-379 | 42.5 | 24.5 | Bulgaria | Europe |
| CGN15705^a^ | TKI-382 | 41.8 | 44.5 | Georgia | West Asia |
| CGN15716^a^ | TKI-383 | 43.3 | 46.0 | Russia | West Asia |
| CGN19047^a^ | TKI-391 | 43.0 | 11.6 | Italy | Europe |
| CGN20697^a^ | TKI-395 | 41.6 | 69.8 | Uzbekistan | Central Asia |

^a^ *L. saligna* accessions used for leaf imaging (Supplemental Figure 6).

^b^ Obtained from the French botanical garden: Jardin Botanique de Nantes.

^c^ According to the CGN passport, CGN05327 was collected at “Caca De La Selia, Gerona”. The coordinate of Girona (41.9° N, 2.8° E) was used to indicate its Spanish origin in Figure 1.

**Supplemental Table 13** Summary of *L. saligna* re-sequencing

| **Accession** | **SSD** | **Depth** | **Size (Gb)** |
| --- | --- | --- | --- |
| CGN05271 | TKI-342 | 13.29 | 30.19 |
| CGN05282 | TKI-343 | 7.14 | 16.22 |
| CGN05301 | TKI-344 | 7.85 | 17.82 |
| CGN05304 | TKI-345 | 6.61 | 15.02 |
| CGN05318 | TKI-355 | 6.57 | 14.91 |
| CGN05327 | TKI-364 | 6.34 | 14.41 |
| CGN05330 | TKI-366 | 7.89 | 17.91 |
| CGN05947 | TKI-369 | 8.51 | 19.32 |
| CGN13326 | TKI-374 | 7.85 | 17.82 |
| CGN13330 | TKI-376 | 4.30 | 9.78 |
| CGN13375 | TKI-379 | 7.14 | 16.22 |
| CGN15705 | TKI-382 | 9.63 | 21.87 |
| CGN15716 | TKI-383 | 5.63 | 12.79 |
| CGN19047 | TKI-391 | 3.89 | 8.83 |
| CGN20697 | TKI-395 | 6.12 | 13.90 |

**Supplemental Table 14** Summary of filtered SNPs for 15 SSD lines against *L. saligna* reference genome (v4)

| **Accession** | **SSD** | **Missing**  **site** |  | **Reference**  **site** |  | **SNP site*** | | |  | **Total site** |
| --- | --- | --- | --- | --- | --- | --- | --- | --- | --- | --- |
|  |  |  |  |  |  | **Homozygous** |  | **Heterozygous** |  |  |
| CGN05271 | TKI-342 | 30,705 |  | 4,606,002 |  | 439,611 |  | 94,161 |  | 5,170,479 |
| CGN05282 | TKI-343 | 246,090 |  | 3,639,847 |  | 1,207,601 |  | 76,941 |  | 5,170,479 |
| CGN05301 | TKI-344 | 228,143 |  | 4,544,197 |  | 353,074 |  | 45,065 |  | 5,170,479 |
| CGN05304 | TKI-345 | 385,617 |  | 3,688,012 |  | 1,036,280 |  | 60,570 |  | 5,170,479 |
| CGN05318 | TKI-355 | 395,705 |  | 3,682,362 |  | 1,036,873 |  | 55,539 |  | 5,170,479 |
| CGN05327 | TKI-364 | 209,841 |  | 4,770,289 |  | 162,595 |  | 27,754 |  | 5,170,479 |
| CGN05330 | TKI-366 | 222,906 |  | 3,601,421 |  | 1,267,081 |  | 79,071 |  | 5,170,479 |
| CGN05947 | TKI-369 | 275,558 |  | 3,658,342 |  | 1,165,367 |  | 71,212 |  | 5,170,479 |
| CGN13326 | TKI-374 | 226,246 |  | 4,463,796 |  | 432,166 |  | 48,271 |  | 5,170,479 |
| CGN13330 | TKI-376 | 380,770 |  | 4,143,970 |  | 599,834 |  | 45,905 |  | 5,170,479 |
| CGN13375 | TKI-379 | 209,840 |  | 4,428,077 |  | 477,701 |  | 54,861 |  | 5,170,479 |
| CGN15705 | TKI-382 | 106,396 |  | 3,912,788 |  | 914,160 |  | 237,135 |  | 5,170,479 |
| CGN15716 | TKI-383 | 218,143 |  | 4,188,005 |  | 696,392 |  | 67,939 |  | 5,170,479 |
| CGN19047 | TKI-391 | 333,845 |  | 4,479,632 |  | 322,485 |  | 34,517 |  | 5,170,479 |
| CGN20697 | TKI-395 | 177,320 |  | 4,286,821 |  | 637,011 |  | 69,327 |  | 5,170,479 |

*Biallelic SNPs after hard-filtering and genotype filtering (missing rate and minor allele frequency).

**Supplemental Table 15** Annotation of filtered SNPs by SnpEff

| **SNP annotation** |  |  |
| --- | --- | --- |
| SNP prediction overview | Genome | *Lactuca saligna* assembly v4 |
|  | Number of variants | 5,170,479 |
|  | Number of effects | 6,552,355 |
|  | Genome total length | 2,165,762,035 |
| *n* of effects by functional class | Type | Count |
|  | MISSENSE | 59,930 (0.91%) |
|  | NONSENSE | 2,424 (0.04%) |
|  | SILENT | 47,879 (0.73%) |
|  | Missense / Silent | 1.25 |
| *n* of effects by region | Type | Count |
|  | DOWNSTREAM | 677,381 (10.34%) |
|  | EXON | 112,986 (1.72%) |
|  | INTERGENIC | 4,887,566 (74.59%) |
|  | INTRON | 162,740 (2.48%) |
|  | SPLICE_SITE_ACCEPTOR | 593 (0.01%) |
|  | SPLICE_SITE_DONOR | 378 (0.01%) |
|  | SPLICE_SITE_REGION | 7,803 (0.12%) |
|  | TRANSCRIPT | 4 (0%) |
|  | UPSTREAM | 678,339 (10.35%) |
|  | UTR_3_PRIME | 16,279 (0.25%) |
|  | UTR_5_PRIME | 8,286 (0.13%) |
| *n* of effects by AA change | Type | count |
|  | synonymous | 51,518 (0.79%) |
|  | nonsynonymous | 59,576 (0.91%) |

**Supplemental Table 16** Chromosomal position and gene count of inversions

| **Chromosome** | **Species** | **Start (Mb)** | **End (Mb)** | **Length (Mb)** | ***n* genes** | **density** |
| --- | --- | --- | --- | --- | --- | --- |
| Chr5 | *L. saligna* | 159.89 | 208.66 | 48.77 | 1,232 | 25.26 |
|  | *L. sativa* | 220.85 | 280.15 | 59.30 | 1,242 | 20.94 |
| Chr8 | *L. saligna* | 59.25 | 103.38 | 44.13 | 1,101 | 24.95 |
|  | *L. sativa* | 79.87 | 136.12 | 56.25 | 1,115 | 19.82 |

**Supplemental Table 17** Genes at the border of putative breaking regions

| **New_id** | **Original_id** | **Orientation** | **Start (bp)** | **End (bp)** |
| --- | --- | --- | --- | --- |
| Sal_IVT8_1 | Lsal_1_v1_gn_8_00001933 | - | 59,109,801 | 59,113,498 |
| Sal_IVT8_2 | Lsal_1_v1_gn_8_00001940 | + | 59,247,351 | 59,249,214 |
| Sal_IVT8_3 | Lsal_1_v1_gn_8_00003173 | - | 103,372,189 | 103,375,131 |
| Sal_IVT8_4 | Lsal_1_v1_gn_8_00003174 | + | 103,451,201 | 103,453,978 |
| Sat_IVT8_1 | Lsat_1_v5_gn_8_57181 | - | 79,537,339 | 79,538,753 |
| Sat_IVT8_2 | Lsat_1_v5_gn_8_94501 | - | 136,118,827 | 136,122,721 |
| Sat_IVT8_3 | Lsat_1_v5_gn_8_58161 | + | 79,867,090 | 79,870,337 |
| Sat_IVT8_4 | Lsat_1_v5_gn_8_94080 | + | 135,454,063 | 135,456,841 |
| Sal_IVT5_1 | Lsal_1_v1_gn_5_00002695 | - | 159,858,509 | 159,860,771 |
| Sal_IVT5_2 | Lsal_1_v1_gn_5_00002696 | - | 159,885,875 | 159,887,024 |
| Sal_IVT5_3 | Lsal_1_v1_gn_5_00004098 | - | 208,657,513 | 208,657,973 |
| Sal_IVT5_4 | Lsal_1_v1_gn_5_00004099 | + | 208,716,880 | 208,719,558 |
| Sat_IVT5_1 | Lsat_1_v5_gn_5_102860 | - | 220,795,208 | 220,797,417 |
| Sat_IVT5_2 | Lsat_1_v5_gn_5_147260 | + | 280,152,608 | 280,154,091 |
| Sat_IVT5_3 | Lsat_1_v5_gn_5_102841 | + | 220,854,303 | 220,854,799 |
| Sat_IVT5_4 | Lsat_1_v5_gn_5_147300 | + | 280,179,374 | 280,181,709 |

**Supplemental Table 18** Genes flanking interspecific inversions between *L. saligna* and *L. sativa*

| ***L. saligna*** | ***L. sativa*** | **TAIR homolog** | **Biological process** |
| --- | --- | --- | --- |
| Sal_IVT5_1 | Sat_IVT5_1 | AT4G28950.1 | Regulation of cell shape |
| Sal_IVT5_2 | Sat_IVT5_2 | AT1G65980.1 | Cell redox homeostasis |
| Sal_IVT5_3 | Sat_IVT5_3 | AT2G21820.1 | Seed development |
| Sal_IVT5_4 | Sat_IVT5_4 | AT3G05580.1 | Cell wall integrity |
| Sal_IVT8_1 | Sat_IVT8_1 | AT3G63310.1 | Negative regulation of cell death |
| Sal_IVT8_2 | Sat_IVT8_2 | AT3G63410.1 | Vitamin E biosynthetic process |
| Sal_IVT8_3 | Sat_IVT8_3 | AT5G19990.3 | Protein catabolic process |
| Sal_IVT8_4 | Sat_IVT8_4 | AT1G52780.1 | Signal transduction |

**Supplemental Table 19.** ID conversion of genes flanking interspecific inversion

| ***L. saligna* new id** | ***L. saligna* original id** | ***L. sativa* new ID** | ***L. sativa* original ID** |
| --- | --- | --- | --- |
| Sal_IVT5_1 | Lsal_1_v1_gn_5_00002695 | Sat_IVT5_1 | Lsat_1_v5_gn_5_102860 |
| Sal_IVT5_2 | Lsal_1_v1_gn_5_00002696 | Sat_IVT5_2 | Lsat_1_v5_gn_5_147260 |
| Sal_IVT5_3 | Lsal_1_v1_gn_5_00004098 | Sat_IVT5_3 | Lsat_1_v5_gn_5_102841 |
| Sal_IVT5_4 | Lsal_1_v1_gn_5_00004099 | Sat_IVT5_4 | Lsat_1_v5_gn_5_147300 |
| Sal_IVT8_1 | Lsal_1_v1_gn_8_00001933 | Sat_IVT8_1 | Lsat_1_v5_gn_8_57181 |
| Sal_IVT8_2 | Lsal_1_v1_gn_8_00001940 | Sat_IVT8_2 | Lsat_1_v5_gn_8_94501 |
| Sal_IVT8_3 | Lsal_1_v1_gn_8_00003173 | Sat_IVT8_3 | Lsat_1_v5_gn_8_58161 |
| Sal_IVT8_4 | Lsal_1_v1_gn_8_00003174 | Sat_IVT8_4 | Lsat_1_v5_gn_8_94080 |

**Supplemental Table 20** *NLR* gene number in *L. saligna* and *L. sativa*

| **Type** | **Structure** | ***n* genes** | |
| --- | --- | --- | --- |
|  |  | ***L. saligna* v4** | ***L. sativa* v8** |
| **CNL type^a^** | CNL | 72 | 92 |
|  | CN | 11 | 8 |
|  | NcL | 34 | 50 |
|  | Nc | 22 | 12 |
|  | Total | 139 | 162^b^ |
| **TNL type** | TNL | 133 | 170 |
|  | TN | 4 | 2 |
|  | NtL | 38 | 30 |
|  | Nt | 9 | 0 |
|  | Total | 184 | 202 |
| **Total** | | 323 | 364 |

^a^ RPW8 and Rx_N type of CNL included in this study.

^b^ Lsat_1_v5_gn_9_123601 annotated as CNL in this study.

**Supplemental Table 21** Genomic position of Major Resistance Clusters (MRCs) in *Lactuca saligna*

| **MRCs** | **Start (bp)** | **End (bp)** | **Size (Mb)** | ***n* *NLR*s** |
| --- | --- | --- | --- | --- |
| lsal-MRC1 | 60,999,383 | 106,534,200 | 46 | 75 |
| lsal-MRC2 | 1,654,628 | 47,280,033 | 46 | 55 |
| lsal-MRC3 | 130,826,820 | 156,658,407 | 26 | 16 |
| lsal-MRC4 | 226,059,448 | 237,370,839 | 11 | 13 |
| lsal-MRC5 | 56,398,680 | 208,199,924 | 152 | 8 |
| lsal-MRC8A | 2,370,358 | 15,793,241 | 16 | 3 |
| lsal-MRC8B | 50,240,655 | 51,336,481 | 1 | 16 |
| lsal-MRC8C | 131,831,946 | 149,404,502 | 18 | 18 |
| lsal-MRC9A | 21,304,606 | 76,924,983 | 56 | 8 |
| lsal-MRC9B | 115,677,911 | 136,622,881 | 21 | 16 |
| lsal-MRC9C | 157,169,118 | 160,490,408 | 3 | 8 |

**Supplemental Table 22** *NLR* clusters (NCs) in the *Lactuca saligna* v4 assembly

| **NCs** | **Start (bp)** | **End (bp)** | **Size (Mb)** | ***n* *NLR*s** | **RGC family** |
| --- | --- | --- | --- | --- | --- |
| NC4 | 38,550,568 | 40,684,360 | 2.1 | 10 | RGC20 |
| NC7 | 44,015,670 | 44,478,820 | 0.5 | 11 | RGC14, RGC40 |

**Supplemental Table 23** Number of *NLR*s per RGC family in *L. sativa* and *L. saligna*

|  | **Singletons** | |  |  |  | **Multigene family** | |  |
| --- | --- | --- | --- | --- | --- | --- | --- | --- |
| **RGC** | ***L. sativa*** | ***L. saligna*** | **Type** |  | **RGC** | ***L. sativa*** | ***L. saligna*** | **Type** |
| RGC10 | 1 | 1 | CNL |  | RGC18^a^ | 28 | 25 | CNL |
| RGC22 | 1 | 0 | CNL |  | RGC21^a,b,c^ | 29 | 23 | CNL |
| RGC24 | 1 | 1 | CNL |  | RGC1^a,c^ | 21 | 16 | CNL |
| RGC26 | 1 | 0 | CNL |  | RGC2^b^ | 21 | 22 | CNL |
| RGC28^a^ | 1 | 1 | CNL |  | RGC41^b^ | 13 | 8 | CNL |
| RGC3 | 1 | 1 | CNL |  | RGC9^c^ | 12 | 7 | CNL |
| RGC30 | 1 | 0 | CNL |  | RGC17 | 6 | 4 | CNL |
| RGC31 | 1 | 1 | CNL |  | RGC29^a,c^ | 5 | 13 | CNL |
| RGC39 | 1 | 2 | CNL |  | RGC7 | 3 | 2 | CNL |
| RGC44 | 1 | 1 | CNL |  | RGC27 | 3 | 0 | CNL |
| RGC5^a^ | 1 | 1 | CNL |  | RGC23 | 0 | 2 | CNL |
| RGC13^a^ | 1 | 1 | TNL |  | RGC25^a^ | 2 | 2 | CNL |
| RGC35 | 1 | 1 | TNL |  | RGC32 | 2 | 1 | CNL |
| RGC36 | 1 | 0 | TNL |  | RGC33 | 2 | 3 | CNL |
| RGC37^a^ | 1 | 0 | TNL |  | RGC42^a^ | 1 | 1 | CNL |
| RGC38 | 1 | 0 | TNL |  | RGC6 | 2 | 1 | CNL |
|  |  |  |  |  | RGC16^a-c^ | 52 | 65 | TNL |
|  |  |  |  |  | RGC12^a-b^ | 54 | 50 | TNL |
|  |  |  |  |  | RGC4^a,c^ | 40 | 27 | TNL |
|  |  |  |  |  | RGC14^b,c^ | 15 | 10 | TNL |
|  |  |  |  |  | RGC40^b^ | 10 | 6 | TNL |
|  |  |  |  |  | RGC15^a^ | 12 | 10 | TNL |
|  |  |  |  |  | RGC8^a,c^ | 7 | 1 | TNL |
|  |  |  |  |  | RGC20^a,c^ | 5 | 9 | TNL |
|  |  |  |  |  | RGC34^a^ | 3 | 4 | TNL |

^a^ RGCs containing complete *NLR* (supported by Supplemental Figure 9 and Supplemental Data 5C-D).

^b^ Previous RGC classification updated by phylogeny tree in this study (see Supplemental Dataset 5E) (Christopoulou *et al.*, 2015).

^c^ RGC families contracted or expanded compared to *L. sativa* (count difference ≥ 3 and change ≥ 50%).

**Supplemental Table 24** Pfam HMM motifs used for *RLK* classification

| **Domain** | **Pfam** | **Description** |
| --- | --- | --- |
| B_lectin | PF01453.26 | D-mannose binding lectin |
| EGF_CA | PF07645.17 | Calcium-binding EGF domain |
| GDPD | PF03009.19 | Glycerophosphoryl diester phosphodiesterase family |
| GUB_WAK_bind | PF13947.8 | Wall-associated receptor kinase galacturonan-binding |
| Lectin_C | PF00059.23 | Lectin C-type domain |
| Lectin_legB | PF00139.21 | Legume lectin domain |
| LRR_1 | PF00560.35 | Leucine Rich Repeat |
| LRR_2 | PF07723.15 | Leucine Rich Repeat |
| LRR_3 | PF07725.14 | Leucine Rich Repeat |
| LRR_4 | PF12799.9 | Leucine Rich Repeat |
| LRR_5 | PF13306.8 | Leucine Rich Repeat |
| LRR_6 | PF13516.8 | Leucine Rich Repeat |
| LRR_8 | PF13855.8 | Leucine Rich Repeat |
| LRR_9 | PF14580.8 | Leucine Rich Repeat |
| LTP_2 | PF14368.8 | Probable lipid transfer |
| LysM | PF01476.22 | LysM domain |
| Malectin | PF11721.10 | Malectin domain |
| Malectin_like | PF12819.9 | Malectin-like domain |
| PAN_1 | PF00024.28 | PAN domain |
| PAN_2 | PF08276.13 | PAN-like domain |
| PAN_4 | PF14295.8 | PAN domain |
| PRIMA1 | PF16101.7 | Proline-rich membrane anchor 1 |
| RCC1_2 | PF13540.8 | Regulator of chromosome condensation (RCC1) repeat |
| RVT_2 | PF07727.16 | Reverse transcriptase (RNA-dependent DNA polymerase) |
| Stress-antifung | PF01657.19 | Salt stress response/antifungal |
| SWIM | PF04434.19 | SWIM zinc finger |
| S_locus_glycop | PF00954.22 | S-locus glycoprotein domain |
| Thaumatin | PF00314.19 | Thaumatin family |
| WAK | PF08488.13 | Wall-associated kinase |
| WAK_assoc | PF14380.8 | Wall-associated receptor kinase C-terminal |

**Supplemental Table 25** Hybrid incompatibility (HI) regions mapping in *L. saligna* v4 assembly

| **qseqid** | **sseqid** | **pident** | **length** | **mismatch** | **gapopen** | **qstart** | **qend** | **sstart** | **send** | **evalue** | **bitscore** |
| --- | --- | --- | --- | --- | --- | --- | --- | --- | --- | --- | --- |
| lg_5_122144297 | chr5 | 99.33 | 150 | 0 | 1 | 1 | 150 | 90,640,672 | 90,640,524 | 8.00E-71 | 270 |
| lg_5_301038425 | ch5 | 99.33 | 150 | 0 | 1 | 1 | 150 | 227,563,284 | 227,563,136 | 8.00E-71 | 270 |
| lg_8_046511832 | chr8 | 99.33 | 150 | 0 | 1 | 1 | 150 | 33,149,407 | 33,149,555 | 8.00E-71 | 270 |
| lg_8_177227692 | chr8 | 99.33 | 150 | 0 | 1 | 1 | 150 | 138,074,032 | 138,074,180 | 8.00E-71 | 270 |

**Supplemental Table 26** NHR intervals and *R* gene locus mapping in *L. saligna* v4 assembly

| **qseqid** | **sseqid** | **pident** | **length** | **mismatch** | **gapopen** | **qstart** | **qend** | **sstart** | **send** | **evalue** | **bitscore** |
| --- | --- | --- | --- | --- | --- | --- | --- | --- | --- | --- | --- |
| lg_4_308156420^a^ | chr4 | 99.33 | 150 | 0 | 1 | 1 | 150 | 241,765,106 | 241,765,254 | 8.00E-71 | 270 |
| lg_4_375657989^a^ | chr4 | 99.33 | 150 | 0 | 1 | 1 | 150 | 278,753,273 | 278,753,125 | 8.00E-71 | 270 |
| lg_7_045594790^a^ | chr7 | 99.33 | 150 | 0 | 1 | 1 | 150 | 28,137,366 | 28,137,514 | 8.00E-71 | 270 |
| lg_7_078029918^a^ | chr7 | 99.33 | 150 | 0 | 1 | 1 | 150 | 55,583,560 | 55,583,708 | 8.00E-71 | 270 |
| lg_7_126358820^a^ | chr7 | 99.33 | 150 | 0 | 1 | 1 | 150 | 97,838,993 | 97,838,845 | 8.00E-71 | 270 |
| lg_7_158098396^a^ | chr7 | 99.33 | 150 | 0 | 1 | 1 | 150 | 124,712,197 | 124,712,345 | 8.00E-71 | 270 |
| lg_8_031811145^a^ | chr8 | 99.33 | 150 | 0 | 1 | 1 | 150 | 22,343,829 | 22,343,977 | 8.00E-71 | 270 |
| lg_8_052278785^a^ | chr8 | 99.33 | 150 | 0 | 1 | 1 | 150 | 38,626,898 | 38,627,046 | 8.00E-71 | 270 |
| CLSM16923^b^ | chr2 | 97.3 | 890 | 24 | 0 | 1 | 890 | 10,479,583 | 10,480,472 | 0 | 1511 |

^a^ NHR interval markers (Giesbers *et al.*, 2018).

^b^ Candidate *R* gene locus determining BLR31 effector recognition (Giesbers *et al.*, 2017).

**Supplemental Table 27** Number of differentially expressed genes in *L. saligna* upon *Bremia* infection

| **Time point** | **Up-regulated** | **Down-regulated** | **Total DEGs** |
| --- | --- | --- | --- |
| 8 hpi | 1222 | 46 | 1268 |
| 24 hpi | 1362 | 326 | 1688 |

**Supplemental Table 28** DEGs related to oomycete responses in *Bremia*-infected *L. saligna*

| ***Lactuca saligna* gene^a^** | **TAIR homolog** | **Other name** | **Description** |
| --- | --- | --- | --- |
| Lsal_1_v1_gn_1_00001954 | AT3G11820 | PEN1 | PEN1 mediates plant immune response: vesicle trafficking in preinvasion contributes to non-host resistance in Arabidopsis |
| Lsal_1_v1_gn_2_00001031 | AT2G34930 | - | disease resistance family protein / LRR family protein |
| Lsal_1_v1_gn_2_00003439 | AT3G11340 | UGT76B1 | a uridine diphosphate-dependent glucosyltransferase that modulates plant defense via glycosylation of N-hydroxypipecolic acid within salicylic acid (SA) mediated signaling pathway |
| Lsal_1_v1_gn_1_00002454 | AT2G32800 | LecRK-S.2 | L-type lectin receptor kinase S.2 |
| Lsal_1_v1_gn_9_00001330 | AT3G56400 | WRKY70 | Member of WRKY Transcription Factor; Group III. Function as activator of SA-dependent defense genes and a repressor of JA-regulated genes. |
| Lsal_1_v1_gn_8_00004656 | AT1G73805 | SARD1 | Encodes SAR Deficient 1 (SARD1), a key regulator for salicylic acid (SA) synthesis, regulation of systemic acquired resistance (SAR) |
| Lsal_1_v1_gn_9_00004094 | AT5G10530 | LecRK-IX.1 | Concanavalin A-like lectin protein kinase family protein, confers *Phytophthora* (oomycete) resistance |
| Lsal_1_v1_gn_5_00002487 | AT5G20230 | BCB | involved in Aluminum stress resistance, involved in cell death and leaf senescence |
| Lsal_1_v1_gn_1_00001172 | AT1G15520 | ABCG40 | ABC transporter family involved in abscisic acid transport, response to salicylic acid |
| Lsal_1_v1_gn_4_00005445 | AT5G14930 | SAG101 | encodes an acyl hydrolase involved in senescence, interactor of EDS1, positive regulation of defense response to oomycetes, |
| Lsal_1_v1_gn_4_00003154 | AT1G59870 | PEN3 | ATP binding cassette transporter. In infected leaves, the protein concentrated at infection sites. Contributes to nonhost resistance to inappropriate pathogens that enter by direct penetration in a salicylic acid–dependent manner. |
| Lsal_1_v1_gn_9_00004137 | AT4G33430 | BAK1 | Leu-rich receptor Serine/threonine protein kinase, involved in protein phosphorylation, contributes to post-invasive immunity |
| Lsal_1_v1_gn_3_00002264 | AT5G06740 | LecRK-S.5 | L-type lectin receptor kinase S.5 |

^a^ See Supplemental Table 29 for expression values.

**Supplemental Table 29** Expression level of DEGs related to oomycete responses in *Bremia*-infected *L. saligna*

| ***Lactuca saligna* gene model** | **8hpi** | | | | |
| --- | --- | --- | --- | --- | --- |
|  | **log2FC** | ***Bremia*** | | **Mock** | |
|  |  | **Mean^a^** | **SD^b^** | **Mean^a^** | **SD^b^** |
| Lsal_1_v1_gn_1_00001954 | 11.72 | 623.17 | 299.47 | 0.00 | 0.00 |
| Lsal_1_v1_gn_2_00001031 | 6.49 | 16.53 | 1.61 | 0.00 | 0.00 |
| Lsal_1_v1_gn_2_00003439 | 4.42 | 1343.33 | 216.44 | 62.54 | 25.99 |
| Lsal_1_v1_gn_1_00002454 | 3.51 | 925 | 202 | 81 | 32.66 |
| Lsal_1_v1_gn_9_00001330 | 3.31 | 40.98 | 12.26 | 4.20 | 3.39 |
| Lsal_1_v1_gn_8_00004656 | 3.30 | 364.49 | 44.31 | 37.01 | 10.30 |
| Lsal_1_v1_gn_9_00004094 | 3.20 | 18.31 | 5.09 | 1.96 | 0.67 |
| Lsal_1_v1_gn_5_00002487 | 3.15 | 232.62 | 19.75 | 26.04 | 21.34 |
| Lsal_1_v1_gn_1_00001172 | 2.71 | 14464.66 | 3043.38 | 2203.65 | 1015.67 |
| Lsal_1_v1_gn_4_00005445 | 1.99 | 77.33 | 14.39 | 19.44 | 4.23 |
| Lsal_1_v1_gn_4_00003154 | 1.66 | 48886.64 | 7138.47 | 15471.68 | 3761.35 |
| Lsal_1_v1_gn_9_00004137 | 1.18 | 270.96 | 64.74 | 119.46 | 17.18 |
| Lsal_1_v1_gn_3_00002264 | 1.17 | 788.41 | 168.58 | 349.22 | 146.79 |
| ***Lactuca saligna* gene model** | **24hpi** | | | | |
|  | **log2FC** | ***Bremia*** | | **Mock** | |
|  |  | **Mean^a^** | **SD^b^** | **Mean^a^** | **SD^b^** |
| Lsal_1_v1_gn_1_00001954 | NDE^c^ | 113.64 | 90.95 | 0.00 | 0.00 |
| Lsal_1_v1_gn_2_00001031 | 4.88 | 6.79 | 2.19 | 0.00 | 0.00 |
| Lsal_1_v1_gn_2_00003439 | 5.43 | 120.96 | 58.10 | 2.63 | 1.90 |
| Lsal_1_v1_gn_1_00002454 | 3.01 | 675.11 | 350.32 | 83.52 | 29.16 |
| Lsal_1_v1_gn_9_00001330 | 3.04 | 30.60 | 9.50 | 3.86 | 2.73 |
| Lsal_1_v1_gn_8_00004656 | 3.10 | 294.50 | 101.29 | 34.46 | 8.43 |
| Lsal_1_v1_gn_9_00004094 | 3.98 | 73.89 | 26.97 | 4.91 | 3.76 |
| Lsal_1_v1_gn_5_00002487 | 4.57 | 280.31 | 183.23 | 11.80 | 1.54 |
| Lsal_1_v1_gn_1_00001172 | 3.74 | 4376.56 | 1827.60 | 327.32 | 20.60 |
| Lsal_1_v1_gn_4_00005445 | 2.94 | 168.20 | 41.94 | 22.12 | 7.81 |
| Lsal_1_v1_gn_4_00003154 | 1.04 | 13733.13 | 4123.32 | 6674.92 | 393.30 |
| Lsal_1_v1_gn_9_00004137 | 1.70 | 703.60 | 159.73 | 216.60 | 25.56 |
| Lsal_1_v1_gn_3_00002264 | NDE^c^ | 479.72 | 159.87 | 327.87 | 18.70 |

^a^ Average of normalized read count for three biological replicates per treatment.

^b^ Standard deviation for three biological replicate samples.

^c^ Not differentially expressed.

**Supplemental Table 30** Statistics of DEGs in four NHR regions of *L. saligna*

| **Region** | **length (Mb)** | **Up-regulated genes** | | **Highly up-regulated genes (Log^2^FC > 3)** | | **Up-regulated *RLK*s*/NLR*s** | | | | | |
| --- | --- | --- | --- | --- | --- | --- | --- | --- | --- | --- | --- |
|  |  |  |  |  |  | ***RLK*** | | | ***NLR*** | | |
|  |  | **n** | **Density/Mb** | **n** | **Density/Mb** | **Total** | **n** | **%** | **Total** | **n** | **%** |
| NHR4 | 36.99 | 20 | 0.54 | 3 | 0.08 | 20 | 1 | 5.00 | 1 | 0 | 0.00 |
| NHR7.1 | 27.45 | 14 | 0.51 | 5 | 0.18 | 2 | 0 | 0.00 | 13 | 0 | 0.00 |
| NHR7.2 | 26.87 | 10 | 0.37 | 2 | 0.07 | 2 | 0 | 0.00 | 0 | 0 | 0.00 |
| NHR8 | 16.28 | 25 | 1.54 | 9 | 0.55 | 14 | 7 | 50.00 | 0 | 0 | 0.00 |
| Genome^*^ | 1,745.02 | 1,222 | 0.7 | 242 | 0.14 | 422 | 96 | 22.70 | 280 | 13 | 4.60 |

* Excluding chromosome 0

**Supplemental Table 31** Candidate genes in NHR8 involved in *L. saligna* nonhost resistance

| **Candidate genes on NHR8** | ***RLK* family** | ***A. thaliana* homolog** | **8hpi** | | | | |
| --- | --- | --- | --- | --- | --- | --- | --- |
|  |  |  | **log2FC** | ***Bremia*** | | **Mock** | |
|  |  |  |  | **Mean^a^** | **SD^b^** | **Mean^a^** | **SD^b^** |
| Lsal_1_v1_gn_8_00000873 | LysM-RK | AT2G33580.1 | 1.60 | 1479.72 | 273.58 | 486.64 | 67.7 |
| Lsal_1_v1_gn_8_00001149 | WAK | AT1G21270.1 | 3.43 | 251.79 | 125.72 | 23.27 | 12.52 |
| Lsal_1_v1_gn_8_00001150 | WAK | AT1G21270.1 | 5.42 | 7.93 | 3.82 | 0 | 0 |
| Lsal_1_v1_gn_8_00001155 | WAK | AT1G21270.1 | 3.75 | 466.14 | 174.77 | 34.71 | 11.33 |
| Lsal_1_v1_gn_8_00001221 | G-LecRK | AT5G60900.1 | 2.85 | 36.1 | 8.22 | 5 | 1.73 |
| Lsal_1_v1_gn_8_00001226 | G-LecRK | AT5G60900.1 | 1.05 | 128.34 | 22.4 | 61.88 | 7.73 |
| Lsal_1_v1_gn_8_00001239 | NA^c^ | AT5G37490.1 | 3.90 | 254.25 | 67.4 | 16.91 | 11.08 |
| Lsal_1_v1_gn_8_00001242 | G-LecRK | AT5G24080.2 | 1.79 | 1724.15 | 171.93 | 498.64 | 110.18 |

^a^ Average of normalized read count for three biological replicates per treatment.

^b^ Standard deviation based on three biological replicates.

^c^ Encoding a plant U-box type E3 ubiquitin ligase (PUB).
