## Supplemental Note for "The genome of *Lactuca saligna*, a wild relative of lettuce, provides insight into non-host resistance to the downy mildew *Bremia lactucae*"

**The presented reference genome sequence is from *L. saligna* accession CGN05327, while we initially intended to sequence accession CGN05271. Below we explain the background of the sample swap and provide evidence on the origin of all data sets regarding these two accessions.**

### Sequenced accession

All *L. saligna* accessions selected for whole-genome sequencing and resequencing were obtained from the lettuce germplasm collection of the Centre for Genetic Resources, The Netherlands (CGN). Accession CGN05327 was collected from Spain (Caca De La Selia, Gerona) and a Single Seed Descendant (SSD) of CGN05327 was used for *de novo* reference genome sequencing and assembly. However, our initial intention was to use the accession CGN05271 for the reference genome assembly. CGN05271 had been crossed in earlier studies with *L. sativa* cv. Olof to develop an F_2_ population and a set of introgression lines (Jeuken & Lindhout, 2001, 2004 ). These materials were used for many in-depth genetic studies on resistance to downy mildew and reproductive barriers (den Boer et al., 2014; Giesbers et al., 2017, 2018, 2019; Jeuken et al., 2002). Importantly, in the above genetic studies of *L. saligna*, the used and reported accession number CGN05271 is true. This finding is based on the fact that the original cross of [*L. saligna* CGN05271 x *L. sativa* Olof] was executed in 1996, before the here identified name-swap incident, and with a different SSD seed batch of CGN05271.

### Origin of the sample swap

A swapping of accession numbers on the seed bags of two different *L. saligna* lines was traced back to 1997. The seeds of aforementioned two accessions (CGN05327 and CGN05271) were both originally obtained from CGN and had independently been crossed to *L. sativa*. The descendants (SSD) of the original cross mothers (i.e., *L. saligna* accessions) were subsequently mislabeled:

- descendent 950240-1A was annotated as CGN05327 while it was CGN05271.
- descendent 950241-1A was annotated as CGN05271 while it was CGN05327.

The mislabeled seed of descendent 950241-1A was used for further regeneration. In the same year (1997), seeds from *L. saligna* accessions CGN05327 and CGN05271 were requested from CGN and also used for regeneration, resulting in regenerations from both the original accessions and the mislabeled accessions. The seeds for the resequencing efforts in this study were requested directly from CGN in 2016. The seeds used by Wei et al. (2021) originate from CGN directly as well.

Between 2015-2017 the seed bag, labeled as ‘CGN05271 pv15257 july 2011’ (SN.Figure 1) was used as the source for plants grown, which were used for RNA sequencing, HiSeq, PacBio sequencing and BioNano sequencing at Wageningen University & Research (WUR) and Hi-C sequencing at UC Davis. Below we present several lines of evidence showing the sequenced genotype, used to construct the reference genome, is CGN05327.

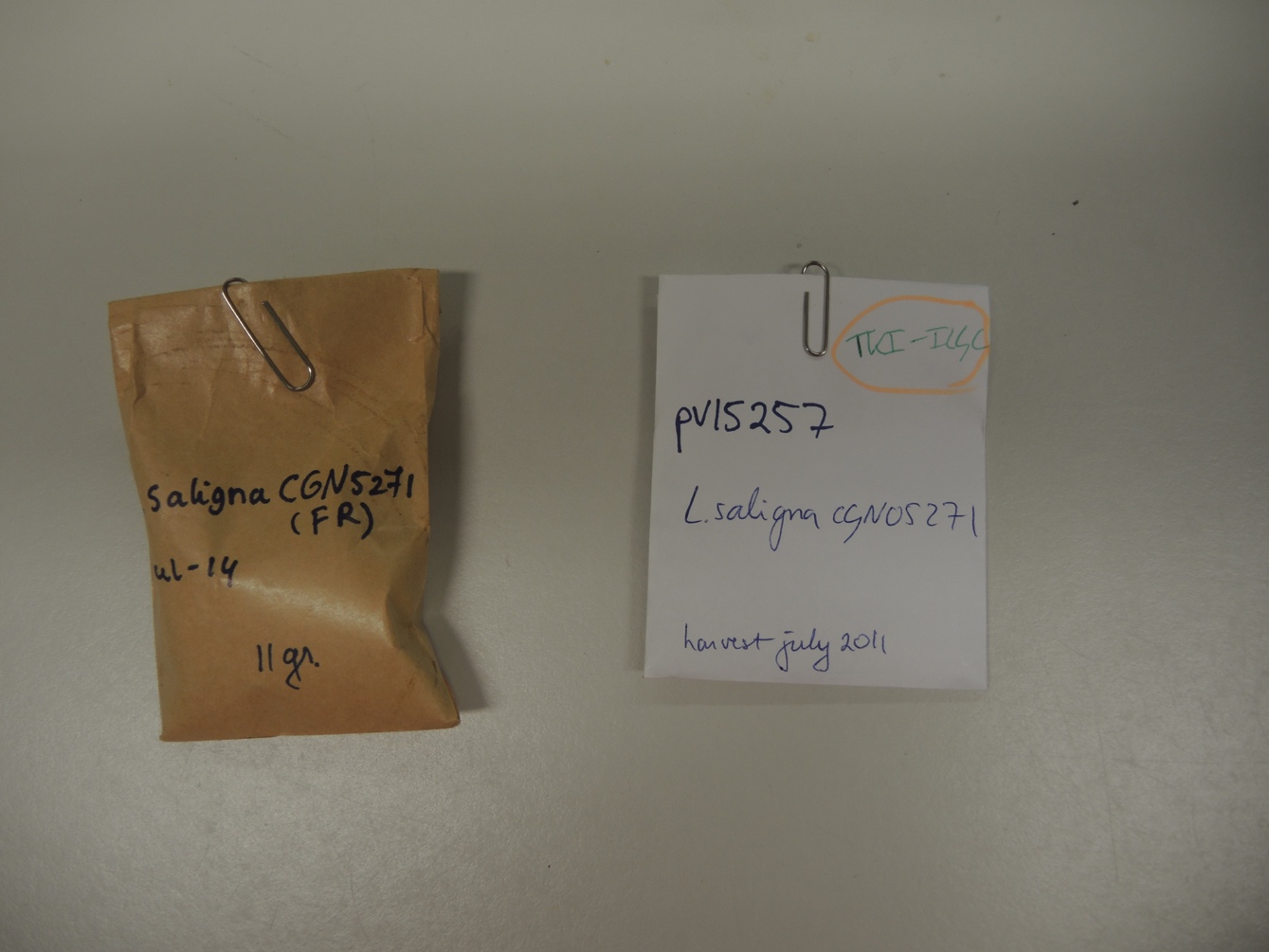

SN.Figure 1: The mislabeled seed bag used for de novo sequencing of *L. saligna*. Although it says ‘CGN05271’, this seed bag contains seeds of CGN05327.

### K-mer analysis of sequencing data confirmed the sample swap

A detailed comparison was performed on Illumina data between sequenced CGN05271 and CGN05327 to confirm this swap. We mainly base our argumentation on k-mer analyses (performed with meryl v1.3 and merqury v1.3 (Rhie et al., 2020)) and further support it with the available variant calling data presented in this paper.

For Illumina genome sequencing, we assume that every region on the genome has an equal chance of being sequenced. Therefore, every region on the genome is expected to have a sequencing depth around the mean of the genome coverage of the entire sequencing data. Thus, when we create a k-mer profile of sequencing data of an inbreeding diploid species (such as *L. saligna*), we expect to see a peak corresponding to the mean genome coverage. Next, a k-mer profile can be used as a unique fingerprint for genetic material: sequencing the same material twice will give the same k-mers, just with a shifted peak depending on genome coverage. If we then compare the k-mer profiles of two genetically distinct accessions, we expect to find unique k-mers under their genome coverage peaks in both k-mer profiles that are absent from the other accession. An important sidenote to make with k-mer profiles of sequencing data is that k-mer profiles always contain a peak at a frequency around 1, corresponding to sequencing artefacts (k-mers that do not occur in the genetic material).

Illustrating this principle, we created this comparison for the resequencing data of CGN05271 (TKI342) and CGN05327 (TKI364) that were sequenced in TKI-100 project (SN.Figure 2A). Clearly, under the peaks corresponding to the mean genome coverage one can identify unique k-mers that are absent from the other accession. This confirms that both accessions are genetically distinct. Next, we compared the resequencing data of CGN05271 with the *de novo* sequencing data for *L. saligna* (which was supposed to be CGN05271 as well), but again we see unique k-mers at the mean genome coverage peak (SN.Figure 2C), indicating these two sequencing datasets originate from genetically distinct material. However, when comparing the resequencing data of CGN05327 with the *de novo* sequencing data for *L. saligna*, we see no unique k-mers at the mean genome coverage peak (SN.Figure 2B), indicating that the *de novo* sequencing data for *L. saligna* corresponds to CGN05327.

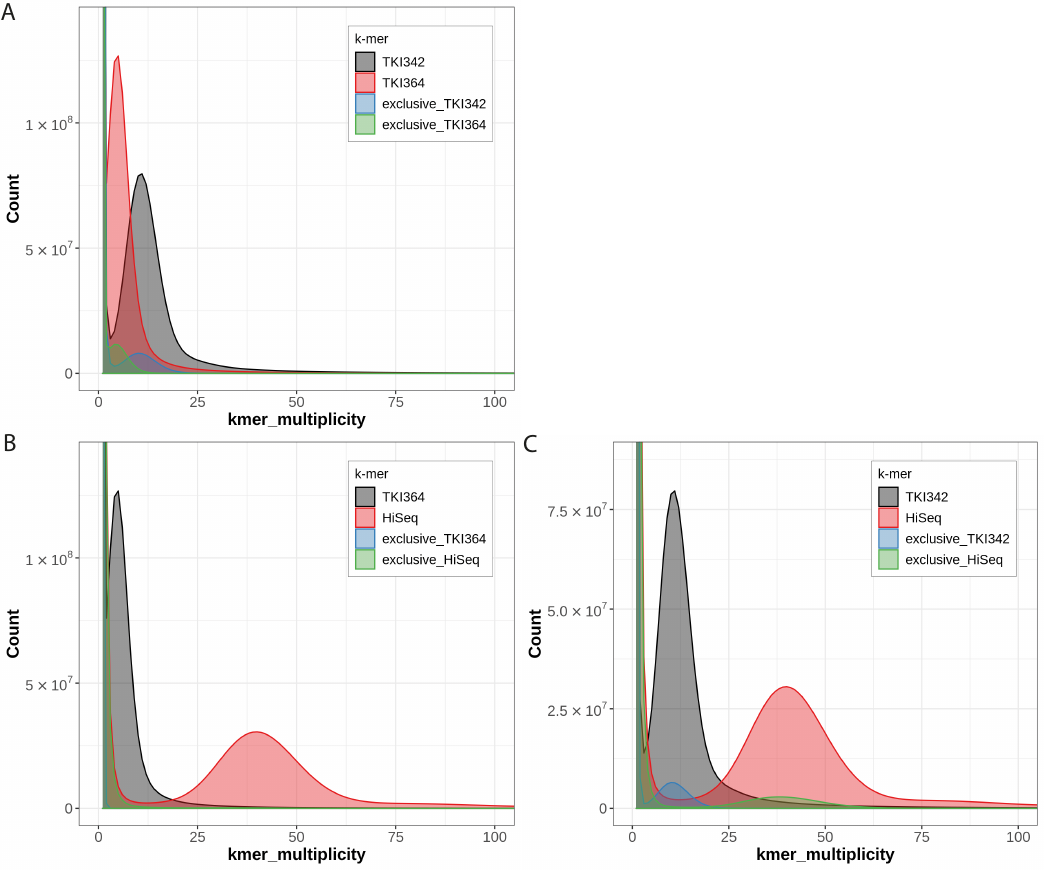

SN.Figure 2: K-mer spectra-cn plot for the pairwise comparisons between TKI342 (CGN05271), TKI364 (CGN05327) and the HiSeq (insert size of 500bp) as sequenced for the *de novo* assembly of *L. saligna*. The black and red lines contain the full k-mer profile for these two sequencing datasets, and the blue and the green lines contain the k-mer profile of the exclusive (unique) content of the indicated line compared to the other.

We reasoned that the number of SNPs identified in these two accessions with variant calling against the *L. saligna* genome should reflect which accession is the one sequenced in the *de novo* sequencing data. Again, CGN05327 looks most like the reference genome as it has the lowest number of SNPs of all accessions used for variant calling (Supplementary Table 14).

Furthermore, we also performed the same k-mer analysis as described above with publicly available resequencing data for both accessions (s333 and s353) (Wei et al., 2021). Again, this analysis proved that the *de novo* sequencing data corresponds to CGN05327 instead of CGN05271 (SN.Table 1).

SN.Table 1: All-vs-all pairwise comparison based on k-mer spectra-cn plots between TKI342 (CGN05271), s333 (CGN05271), TKI364 (CGN05327), s353 (CGN05327) and the HiSeq (insert size of 500bp) as sequenced for the de novo assembly of *L. saligna*. A ‘Y’ indicates similar genetic material, ‘N’ indicates dissimilar genetic material.

| Sample *CGN ID* (dataset) | HiSeq  (*de novo*) | TKI342 CGN05271 (this study) | s333 CGN05271 (Wei et al.) | TKI364 CGN05327 (this study) | s353 CGN05327 (Wei et al.) |
| --- | --- | --- | --- | --- | --- |
| HiSeq  (*de novo*) | - |  |  |  |  |
| TKI342 CGN05271 (this study) | N | - |  |  |  |
| s333 CGN05271 (Wei et al.) | N | Y | - |  |  |
| TKI364 CGN05327 (this study) | Y | N | N | - |  |
| s353 CGN05327 (Wei et al.) | Y | N | N | Y | - |

Finally, making sure that all the sequencing data underlying the *de novo* genome has the same genetic origin, we completed a pairwise comparison of available datasets used here (Illumina HiSeq, MiSeq, 10X, Hi-C) based on k-mer profiles and unique k-mer content. Due to their nature, the PacBio, Bionano and RNA-seq data could not be checked with k-mer profiles but since they originate from the same material, we assume they have the same origin. From these comparisons, we conclude that all sequencing data has the same genetic origin and the *L. saligna* CGN05327 genome assembly is not a chimeric assembly (data not shown since no exclusive content was found at any genome coverage peak).

### Sorting out the datasets

Summarizing, we here provide the origin of each dataset related to this study.

SN.Table 2: The metadata of all sequencing data used for de novo assembly of *L. saligna* CGN05327.

| Type of data | Study origin | Seed bag | Sequenced by | CGN accession |
| --- | --- | --- | --- | --- |
| HiSeq (500bp insert size) | This study | pv15257 | WUR | CGN05327 |
| HiSeq (200bp insert size) | This study | pv15257 | WUR | CGN05327 |
| MiSeq | This study | unknown | WUR | CGN05327 |
| 10X | This study | unknown | WUR | CGN05327 |
| PacBio | This study | pv15257 | WUR | CGN05327 |
| Bionano | This study | pv15257 | WUR | CGN05327 |
| Hi-C | This study | pv15257 | UC Davis | CGN05327 |
| RNA-seq (root) | This study | pv15257 | WUR | CGN05327 |
| RNA-seq (flower) | This study | pv15257 | WUR | CGN05327 |
| RNA-seq (*Bremia* exp.) | This study | pv15257 | WUR | CGN05327 |

SN.Table 3: The metadata of all resequencing data used for the comparisons described in this Supplementary File.

| Sample name | Study origin | Seed origin | Sequenced by | CGN accession |
| --- | --- | --- | --- | --- |
| TKI342 | This study | CGN | WUR | CGN05271 |
| s333 | Wei et al. (2021) | CGN | BGI | CGN05271 |
| TKI364 | This study | CGN | WUR | CGN05327 |
| s353 | Wei et al. (2021) | CGN | BGI | CGN05327 |
